# Bioinformatic discovery of type 11 secretion system (T11SS) cargo across the Proteobacteria

**DOI:** 10.1101/2024.09.26.615229

**Authors:** Alex S. Grossman, Nicholas C. Mucci, Sarah J. Kauffman, Jahirul Rafi, Heidi Goodrich-Blair

## Abstract

<u>T</u>ype <u>11 s</u>ecretion <u>s</u>ystems (T11SS) are broadly distributed among proteobacteria, with more than 3000 T11SS family outer membrane proteins (OMPs) comprising 10 major sequence similarity network (SSN) clusters. Of these, only 7, all from animal-associated cluster 1, have been experimentally verified as secretins of cargo, including adhesins, hemophores, and metal binding proteins. To identify novel cargo of a more diverse set of T11SS, we identified gene families co-occurring in gene neighborhoods with either cluster 1 or marine microbe-associated cluster 3 T11SS OMP genes. We developed bioinformatic controls to ensure perceived co-occurrences are specific to T11SS, and not general to OMPs. We found that both cluster 1 and cluster 3 T11SS OMPs frequently co-occur with single carbon metabolism and nucleotide synthesis pathways, but that only cluster 1 T11SS OMPs had significant co-occurrence with metal and heme pathways, as well as with mobile genetic islands, potentially indicating diversified function of this cluster. Cluster 1 T11SS co-occurrences included 2556 predicted cargo proteins, unified by the presence of a C-terminal β-barrel domain, which fall into 141 predicted UniRef50 clusters and approximately 10 different architectures: 4 similar to known cargo and 6 uncharacterized types. We experimentally demonstrate T11SS-dependent secretion of an uncharacterized cargo type with homology to <u>P</u>lasmin <u>s</u>ensitive <u>p</u>rotein (Pls). Unexpectedly, genes encoding marine cluster 3 T11SS OMPs only rarely co-occurred with the C-terminal β-barrel domain and instead frequently co-occurred with DUF1194-containing genes. Overall, our results show that with sufficiently large-scale and controlled genomic data, T11SS-dependent cargo proteins can be accurately predicted.

**Impact Statement:** We describe a novel method for controlling genomic neighborhood analyses and use this controlled co-occurrence technique to examine two distinct clusters of genes from the recently described type 11 secretion system (T11SS): Cluster 1, predominantly encoded by host-associated microbes and Cluster 3, encoded by marine microbes. We found that both clusters of predicted T11SS frequently co-occur with single carbon metabolism and nucleotide synthesis pathways, but only the host-associated cluster co-occurred with iron uptake pathways. Using these datasets, we predicted 2687 T11SS dependent cargo with approximately ten unique architectures, six of which have not previously been linked to T11SS. Finally, we validate our results by demonstrating T11SS-dependent secretion of a cargo protein with one of the novel architectures, plasmin-sensitive surface protein Pls from *Haemophilus parahaemolyticus*.

**Data Summary:** This paper describes genomic co-occurrences of T11SS OMP sequences originally identified through sequence similarity networking in a previous publication [3] as constituents of similarity clusters, 1 and 3. Accession IDs and assorted metadata for those clusters/networks are listed in Table S1 of that article: https://journals.asm.org/doi/suppl/10.1128/mbio.01956-21/suppl_file/mbio.01956-21-st001.xlsx. All other supporting data, accession IDs, code, and protocols have been provided within this article or through supplementary data files. Additionally, all supplemental data are available on Figshare (https://doi.org/10.6084/m9.figshare.28514069.v2) [1].

**Graphical abstract:** 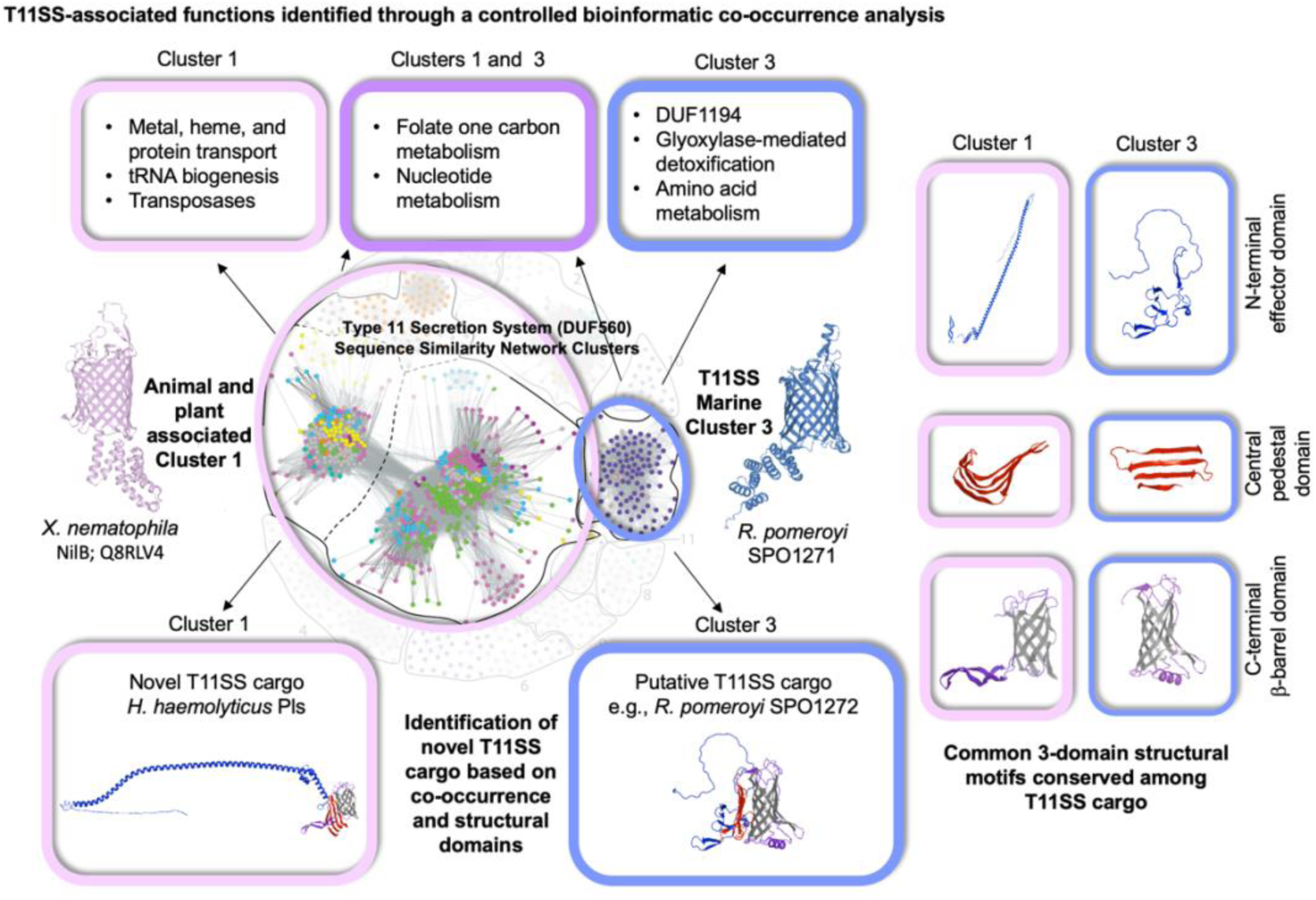

## Introduction

Identifying protein function remains a difficult task in the postgenomic age. As of 12/02/2024 there were 5,366 domains of unknown function (DUF) families, which constitutes around 22.6% of the database [2]. This means roughly one quarter of the Pfam database remains uninvestigated, limiting our understanding of the breadth of biological processes occurring on Earth. Functional characterization of proteins with DUF domains enables their annotation and removal from the DUF list, improving our datasets permanently. *In silico* approaches applied to protein domains are an important component of this process, as they can focus attention on the likely functions of a particular protein family [3]. Recently, a combination of *in silico* and experimental approaches has yielded insight into the DUF560 protein family of outer membrane proteins (OMPs), defining it as the type 11 bacterial secretion system (T11SS), represented by 10 distinct sequence-level clusters [4]. T11SS are outer membrane proteins that translocate either soluble proteins or membrane anchored lipoproteins from the periplasmic space to the extracellular space [4]. Presumably cargo proteins are transported in an unfolded state, since some cargo are dependent on Skp chaperones [5]. T11SS homologs are present among a diverse array of Proteobacteria, including human pathogens in the *Neisseria*, *Haemophilus*, and *Moraxella* genera [6–8]. In the gamma-proteobacterium *Xenorhabdus nematophila*, a T11SS system encoded within the <u>n</u>ematode intestinal localization locus (*nil*) is necessary for colonization of the mutualistic nematode host *Steinernema carpocapsae* [9]. In this system the T11SS, NilB, regulates the surface exposure of a surface lipoprotein, NilC, both of which are necessary for colonization. Other known lipidated T11SS-dependent cargo include well studied pathogenesis factors such as the metal uptake co-receptors transferrin binding protein B (TbpB) and lactoferrin binding protein B (LbpB), as well as factor H binding protein, which binds an immune activation factor [6–8, 10, 11]. These cargo are composed of two types of domains, variable N-terminal effector domains and 8-stranded C-terminal β-barrel domains (such as TbpBBD and Lipoprotein C). T11SS can transport unlipidated, soluble cargo proteins across the outer membrane. For example, T11SS-dependent hemophilin homologs have been characterized from *X. nematophila* (HrpC), *Acinetobacter baumannii* (HphA), and *Haemophilus haemolyticus* (Hpl) [4, 10, 12, 13]. However, the majority of T11SS OMPs do not have a known or predicted cognate cargo. A sequence similarity network analysis suggested that experimentally characterized T11SS only capture a small portion of the functional diversity present in this family [4].

The availability and accessibility of sequencing data have skyrocketed since the first bacterial genome was sequenced in 1995, with ∼315,000 prokaryotic genomes on RefSeq [14]. This tremendous repository of genomic sequence data allows computational biologists to probe these genomes for patterns in genetic co-occurrence, as measured through proximity to a gene of interest. Genes within a common functional pathway often cluster together within a genome. Such patterns can be useful when forming hypotheses about the function of uncharacterized proteins. Gene neighborhood studies analyze the genomic regions upstream and downstream of any given gene of interest with the assumption that nearby genetic partners could illuminate the role(s) of the unknown gene [15].

However, one flaw in the current methodology used to derive co-occurrence lists is the lack of bioinformatic controls capable of removing non-specific co-occurring genes. For example, commonly inherited, deeply ancestral domains, such as transcription and translation machinery, often appear as the top result of a co-occurrence analysis, but do not yield useful information about the specific query gene since their appearance is a coincidence of ubiquity [16]. Also, meaningful results can be obscured by co-occurrence based on structural or other common features present within the query protein, such as those that target the gene product to specific cellular locations, which are expected to interact commonly with the relevant protein trafficking machinery. Herein, we use computational approaches to elucidate potential functional roles played by T11SS-dependent cargo and predict novel cargo. Using these results, we then generated a co-expression system in *Escherichia coli* to experimentally demonstrate the secretion of one novel, bioinformatically predicted cargo family using the <u>pl</u>asmin <u>s</u>ensitive protein (Pls) from *Haemophilus parahaemolyticus* and the type <u>e</u>leven <u>P</u>ls <u>S</u>ecretor (TepS).

## Methods

### Gene neighborhood analysis

The T11SS genome neighborhoods (±6 genes around a T11SS gene) from cluster 1 (1111 queries) and cluster 3 (145 queries) [4] were analyzed. T11SS genome neighborhoods were generated through Rapid ORF Detection & Evaluation Online (RODEO) [17]. As described in the main text, a control was developed to identify and remove non-specific genes from among those flagged as co-inherited or syntenic with T11SS (Fig. 1). The primary control list was generated by extracting all GO-term outer membrane proteins from proteobacteria and filtering for sequences similar in length to the median T11SS length (480 – 482 amino acids in length) and removing all DUF560-containing proteins. For both T11SS cluster analyses a random subset of control sequences were chosen equal to the number of queries being analyzed.

**Figure 1.**
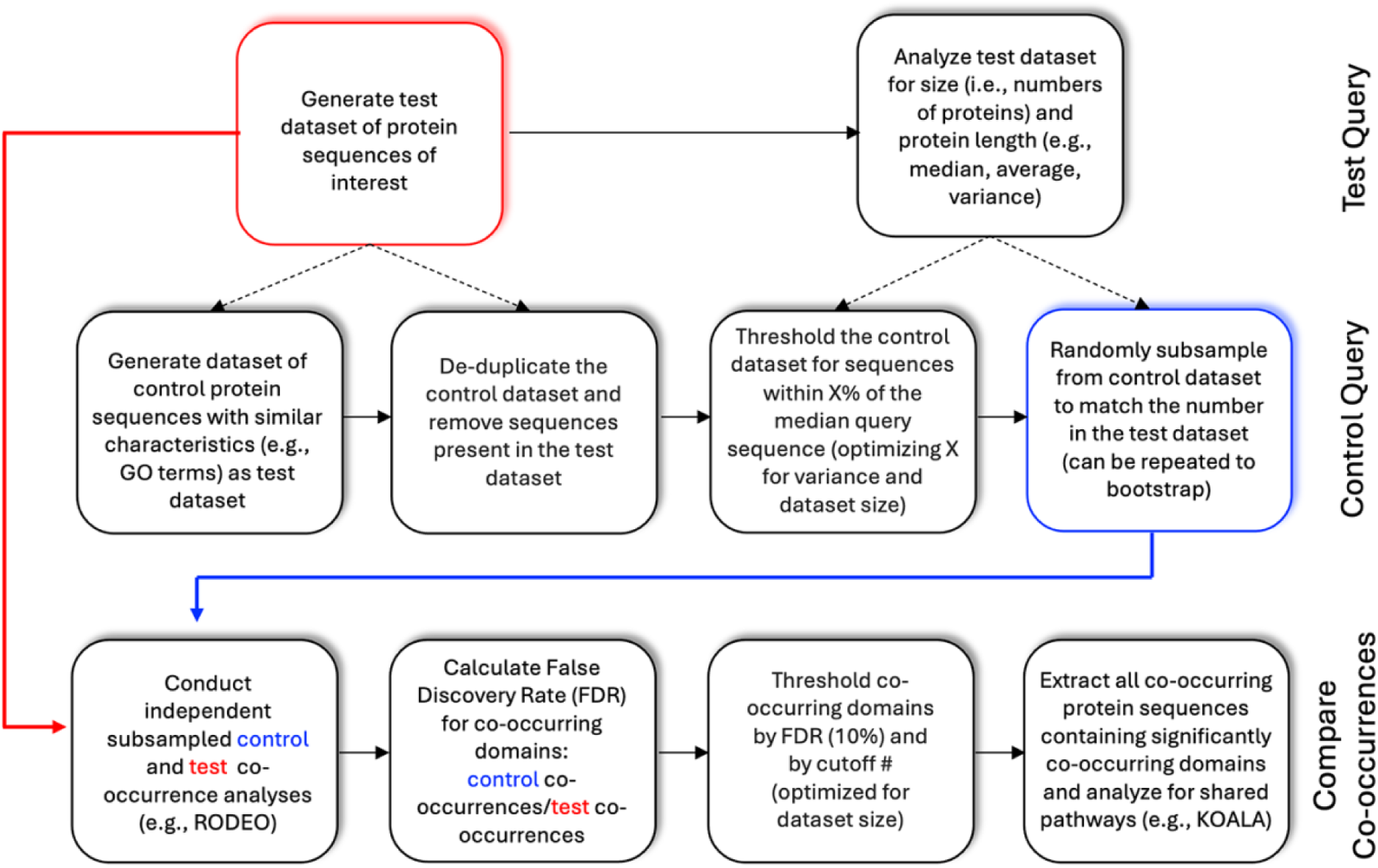
Design for a novel control technique for genomic neighborhood co-occurrence analysis. Logic flowchart to control for nonspecific co-occurrences from a genomic neighborhood analysis. For the analyses presented here, the test dataset queries comprised two different clusters (1 and 3) of T11SS OMPs identified in a published network analysis [4]. Control datasets were developed for each cluster separately. The thresholds for the control datasets were set with X% determined as how many +/-amino acids resulted in the # of proteins that was close to, yet over, each test dataset.

Controlled protein sequences were downloaded from UniProt [18] in February 2021. Co-occurring protein domains were annotated and summed up to determine frequency among queries and controls. False Discovery Rate was calculated by dividing frequency among T11SS queries and control sequences. All domains with and FDR > 0.1 were excluded from analysis. For cluster 1 all domains with fewer than 25 occurrences were excluded. For cluster 3 all domains with fewer than 5 co-occurrences were excluded.

The Joint Genome Institute Integrated Microbial Genomes and Microbiomes platform (https://img.jgi.doe.gov/) [19, 20] was used to examine genomic regions surrounding the T11SS OMPs-encoding genes of select cluster 3 bacteria. Briefly, the “find genes” function was used to identify T11SS OMP homologs from selected representatives of the Roseobacteriaceae and Rhodobacteraceae. The “gene neighborhoods” function was then used to visualize the regions surrounding the T11SS OMP homologs and to identify syntenic loci (for an example, see Supplemental File 3, Neighborhood Alignment Tab). Predicted structures were downloaded as .pdb or .cif files from AlphaFold 2.0 [21, 22] except in one instance (*Amylibacter cionae* CGMCC 1.15880 IEX10_RS12250) for which an AlphaFold 3.0 model [23] was used to predict the structure, and a .pdb file of the top ranked prediction was used for Fig. 8G [21]. Average predicted local distance difference test (pLDDT) values for each predicted structure are provided in Supplementary File 1 (Tab AlphaFold Predicted Structures) Using each .pdb file, structural motifs were highlighted using Protean 3D^®^, Version 17.4.3, DNASTAR, Madison, WI.

### Functional and structural analyses of co-occurring gene neighborhood domains

Protein sequences that contain domains that co-occur with T11SS from the gene neighborhood analysis (Supplemental file 1, Cluster1FDR_Threshold tab) were obtained via UniProt [18]. KofamKOALA [24] and BlastKOALA [25] were used to assess function. KofamKOALA assigns KEGG orthology to user sequence data by HMMER/HMMSEARCH against the KEGG database. BlastKOALA is similar but uses a BLAST search to assign KEGG orthology. Genes were matched to BRITE hierarchies to assess general cellular function and KEGG pathway to assess potential interactions.

To identify potential cargo proteins all downloaded protein sequences were matched to UniRef50 clusters to compress potential homologs [26]. The reference sequences from these UniRef50 clusters where then subjected to a BLAST search against known T11SS cargo β-barrel and handle domains (nematode intestine localization protein (NilC) in *Xenorhabdus nematophila*, transferrin binding protein (TbpB) N-terminal and C-terminal in *Xenorhabdus nematophila*, heme receptor protein (HrpC) in *Xenorhabdus nematophila*, Cobalt receptor protein (CrpC) in *Xenorhabdus cabanillasii*, Hemophilin (Hpl) in *Haemophilus haemolyticus*, hemophilin (HphA) in *Acinetobacter baumanii*, factor H binding protein (fHbp) *Neisseria meningitidis*, and lactoferrin binding protein (LbpB) N-terminal and C-terminal in *Neisseria meningitidis* to search for homologs of T11SS cargo with a low stringency E-value filter of <0.05 in order to find even distantly related homologs, generating a list of 141 potential cargo proteins from cluster 1 and a single potential cargo from cluster 3. MSA alignment was used to group the predicted cargo proteins into distinct architectures. 10 groups emerged, TbpB-like, LbpB-like, Hemophilin-like 1 and 2, fHbp-like, Pls-like, highly disordered N-terminus 1 and 2, *Psychrobacter* surface protein, and *Sphingomonas* surface protein. Structural data predictions for the 9 architectures which contained more than one sequence were generated through PSIPRED [27] for secondary structure and AlphaFold2 [21, 22] for tertiary structure.

### Co-expression of Pls and its cognate T11SS protein TepS

All cultures were grown in glucose minimal media supplemented with 1% LB. Plate based cultures were grown on glucose minimal plates [28]. For plasmid-based expression, chemically competent *Escherichia coli* strain BL21-DE3 (C43) were chosen for ease of transformation and their ability to non-toxically express membrane proteins [29, 30]. Strains of *E coli* were grown at 37°C. Where appropriate media was supplemented with ampicillin at a concentration of 150 μg/ml. Protein expression was induced at the midlog point of bacterial growth via addition of isopropyl β-d-1-thiogalactopyranoside (henceforth IPTG) at a concentration of 0.5mM.

Using sequences from *H. parahaemolyticus* strain NCTC10794, GenScript synthesized genes for Plasmin sensitive surface protein (Pls-STO63610.1) with an attached C-terminal FLAG tag and its genomically associated T11SS OMP (TepS-STO63613.1) with an N-terminal 6xHis tag. Both genes were codon optimized for expression in *E. coli* Bl21DE3. GenScript cloned Pls-FLAG into multiple cloning site 1 of pETDuet-1 and 6xHis-TepS into multiple cloning site 2, resulting in expression plasmids pETDuet-1/Pls-FLAG and pETDuet-1/Pls-FLAG/6xHis-TepS. Expression plasmids were transformed into *E. coli* via electroporation. Strains were grown in defined medium with 150 mg/ml ampicillin [31]. Bacteria were subcultured into 100 ml of broth at an initial optical density at 600 nm (OD_600_) of 0.1, grown for 6 h at 37°C to reach late logarithmic growth, and induced with 0.5 mM IPTG for 1 hour.

### Pls detection assays

Two approaches were taken to monitor localization of Pls: immunoblotting (dot blot and Western) and immunofluorescence and flow cytometry. For the immuno-dot blot, cells were normalized to an OD600 of 1, rinsed 3x, lysed by sonication for 30 sec at ∼500 root mean square volts (V_rms_), and spotted in technical triplicate on nitrocellulose membranes to enumerate total cellular Pls. To monitor extracellular secretion of Pls supernatant was collected from cultures after induction and filtered sterilized to remove all cells. Supernatant samples were ultracentrifuged at 150,000 relative centrifugal force (RCF) for 3 h to separate the soluble fraction from the insoluble component, composed of cell membrane components and membrane vesicles. Previous studies have detected both T11SS OMPs and cargo within membrane vesicles, necessitating their removal from supernatants [4, 32]. The membrane vesicle fraction was suspended in protein sample buffer. 700 μl of each soluble supernatant fraction was precipitated via 10% trichloroacetic acid (TCA) and the resulting pellet was suspended in protein sample buffer [33]. Samples were analyzed by 10% sodium dodecyl sulfate-polyacrylamide gel electrophoresis (SDS-PAGE) and immunoblotting. All dot blots and Western blots were probed with rat anti-FLAG primary antibody and goat anti-rat secondary antibody conjugated to a 680nm fluorophore. Intensities were recorded for FLAG reactive bands. For every supernatant sample, a band from the Coomassie blue-stained gel was used as a loading control to normalize intensities of supernatant samples prior to analysis. Fold change of secretion was determined by dividing the amount of supernatant Pls detected in the Pls/TepS co-expression treatment by the amount detected in the Pls alone treatment. A Tukey’s honestly significant difference (HSD) test was used for comparing fold change of secretion [34].

For immunofluorescence labeling and detection of Pls, overnight cultures of strains were washed thrice with PBS, and resuspended in defined medium at an OD_600_ of 0.05. Cultures were then grown to mid-log phase and one replicate culture of every strain was induced with IPTG at a final concentration of 0.5 mM, while the other culture was left uninduced. All strains were allowed to grow for an additional 1.5 h at 30°C with shaking. Cells were blocked from nonspecific binding using 3% BSA in PBS and incubated with a 1:300 dilution of DYKDDDDK Tag Recombinant Rabbit Monoclonal Antibody Alexa Fluor Plus 488 (Invitrogen) in 0.5% BSA in PBS. Cells were washed thrice in 0.5% BSA in PBS, fixed with 4% formaldehyde in PBS for 15 min, and washed thrice with PBS. Cells were resuspended in PBS and then passaged through a Cytoflex (Cytek NL-3000) and 100,000 events were captured. Visualization and data analyses were performed in FlowJo (Beckton, Dickinson and Company, Franklin Lakes, NJ, USA). Flow cytometry analysis was performed three separate times, and for each experiment three biological replicates were performed, resulting in a total of 9 biological replicates. To monitor total Pls levels expressed, samples from cultures grown for flow experiment were pelleted, washed with PBS, and frozen at −20 for 24 h before being resuspended in PBS and lysed by sonication for 1 min total (5s on, 10s off, 600 V_rms_). After pelleting cellular debris, the protein concentrations of the cell lysate supernatants were determined using the Pierce 660nm Protein Assay Reagent. All samples were normalized and 9 µg of total protein for each treatment were separated by SDS-PAGE gel, followed by Western blot probing with primary anti-DYKDDDDK antibody (63703 Biolegend) at a 1:10,000 dilution and secondary IRDye 680RD Goat anti-Mouse (926-68070) at a 1:10,000 dilution. Blots were imaged using an Odyssey FC.

## Results

### Establishing a control for domain of unknown function (DUF) gene neighborhood co-occurrence analysis

Characterized T11SS-cargo pairs encoded by Proteobacteria typically have conserved roles in host-metal acquisition and immune evasion [4, 8, 10, 12]. Only a small portion of predicted T11SS-cargo pairs has been experimentally characterized, leaving open the possibility that as-yet uncharacterized systems may be participating in diverse symbiotic and metabolic pathways. To gain insights into the potential breadth of biological processes in which T11SS-cargo pairs may function, we collected all T11SS sequences identified as part of the largest T11SS sequence similarity network cluster described in Grossman et al., 2021 (cluster 1, containing 1111 genes) and utilized the Rapid ORF Description & Evaluation Online (RODEO) software to identify all protein domains encoded within their T11SS loci, herein defined as six open reading frames upstream or downstream of the T11SS OMP query genes. Analysis of all loci resulted in a data set of 1504 domains, co-occurring between 2 and 928 times (Supplemental file 1, Cluster1Co tab) [17]. To avoid spurious correlations, we chose to exclude any domains that co-occurred fewer than 25 times (∼2.7% of the maximum frequency co-occurrence). This number was arrived at by generating a histogram of co-occurrence frequency and identifying the inflection point between the rare co-occurring sequences which made up the bulk of the dataset (1331 or 88.5%) and those domains whose co-occurrence frequency was greater than expected randomly (173 or 11.5%) (Fig. S1).

To ensure that the observed co-occurrences were specific to the T11SS protein family and not due to common features of genomic loci encoding outer membrane proteins, we developed and implemented a control process. Briefly, RODEO co-occurrence neighborhoods were generated using randomly selected proteins with biophysical characteristics similar to our DUF560 query proteins such as protein length and outer membrane localization (GO term: outer membrane GO0019867), and then used as a point of comparison for gene neighborhoods from T11SS genes. The control sequences obtained were limited to the Proteobacteria phylum, where T11SS predominantly occur [4]. Due to a left-skewed distribution, the median length of all DUF560 domain containing proteins in Pfam (accessed February 2021) was used to reduce bias from pseudogenes and gene fragments (Fig. S2) [2]. The median size of all predicted T11SS proteins was 481 amino acids, so we extracted all outer membrane ORFs between 480 and 482 amino acids in size. Sequences that appeared in the T11SS database were removed from the control database to ensure unbiased selection. A random subsample of this control database was taken equal to the number of sequences in the original query database (Supplemental file 1, ControlQueries tab) (Fig. 1).

OMP sequences from the subsampled control dataset were then submitted as a query to the genome neighborhood network function of RODEO using the same parameters as the T11SS query list. The output of this control co-occurrence analysis was used to generate a false discovery rate (FDR) for any given domain by dividing frequency of co-occurrence with random OMPs by frequency of co-occurrences with T11SS genes. We then gated the dataset for domains with an FDR less than 0.1 (10%) to exclude non-specific correlations, resulting in a final set of 51 significantly co-occurring domains. (Supplemental file 1, Cluster1FDR_Threshold tab). The most common significant co-occurring domains alongside cluster 1 T11SS OMPs were other DUF560 proteins (928/1111), the TbpBBD β barrel domain (503/1111), the TonB C-terminal domains (364/1111), heme oxygenase (137/1111), and DUF454 (118/1111). Next, all genes that co-occurred with a cluster 1 T11SS gene and possessed one of the identified domains were extracted resulting in 3405 significantly co-occurring genes (Supplemental file 1, Cluster1SigGenes tab). These genes were submitted to KofamKOALA and BlastKOALA in parallel for annotation into molecular pathways and BRITE hierarchies [24, 25].

### Functional assessment of T11SS co-occurring proteins identifies a role of iron uptake, protein export, and single carbon metabolism

We examined the co-occurrence gene set for those with annotations in the curated Kyoto Encyclopedia of Genes and Genomes (KEGG) ortholog (KO) database [24, 35]. A hidden Markov model search using KofamKOALA (HMMER/HMMSEARCH) was used to annotate the genes according to the KO database. The software found matches for 1488 of the 3405 significant co-occurring genes (Supplemental file 2, Cluster1KofamBrite tab). When matched to cellular functions (BRITE hierarchies) the most common categories were transporters (periplasmic TonB, ExbD, Signal peptidase II, TolA, etc.), enzymes (heme oxygenase, formate dehydrogenase, exopolyphosphatase, dihydrofolate reductase, etc.), and tRNA biogenesis (tRNA modifying GTPase, tryptophanyl-tRNA synthase, Ribonuclease P, aminoacyl tRNA synthase) (Fig. 2A). Additionally, cluster 1 T11SS genes were found to significantly co-occur with transposase genes. Of the 1488 matched sequences, 259 had known pathway association, most commonly biosynthesis of secondary metabolites (heme oxygenase, cobalamin-dependent methionine synthase, vitamin K biosynthesis protein MenH, etc.), porphyrin metabolism (heme oxygenase), and protein export (signal peptidase II and YidC insertase) (Fig. 2B) (Supplemental file 2, Cluster1KofamPath tab). The list of pathways also included aminoacyl tRNA biosynthesis (TyrS, TrpS). A high frequency of co-occurrence with transposases and tRNA synthesis genes is characteristic of pathogenicity islands, phage regions, and other mobile genetic elements, since many mobile genetic elements have targeted insertion sites near universally conserved features, such as tRNA genes [36]. Co-occurrence with mobile genetic island signifiers supports our previous observation of horizontal acquisition of a T11SS in *X. nematophila* [4, 9].. Finally, the list included several carbohydrate biosynthesis pathways (hsa00670, one carbon pool by folate; map00230, purine metabolism).

**Figure 2.**
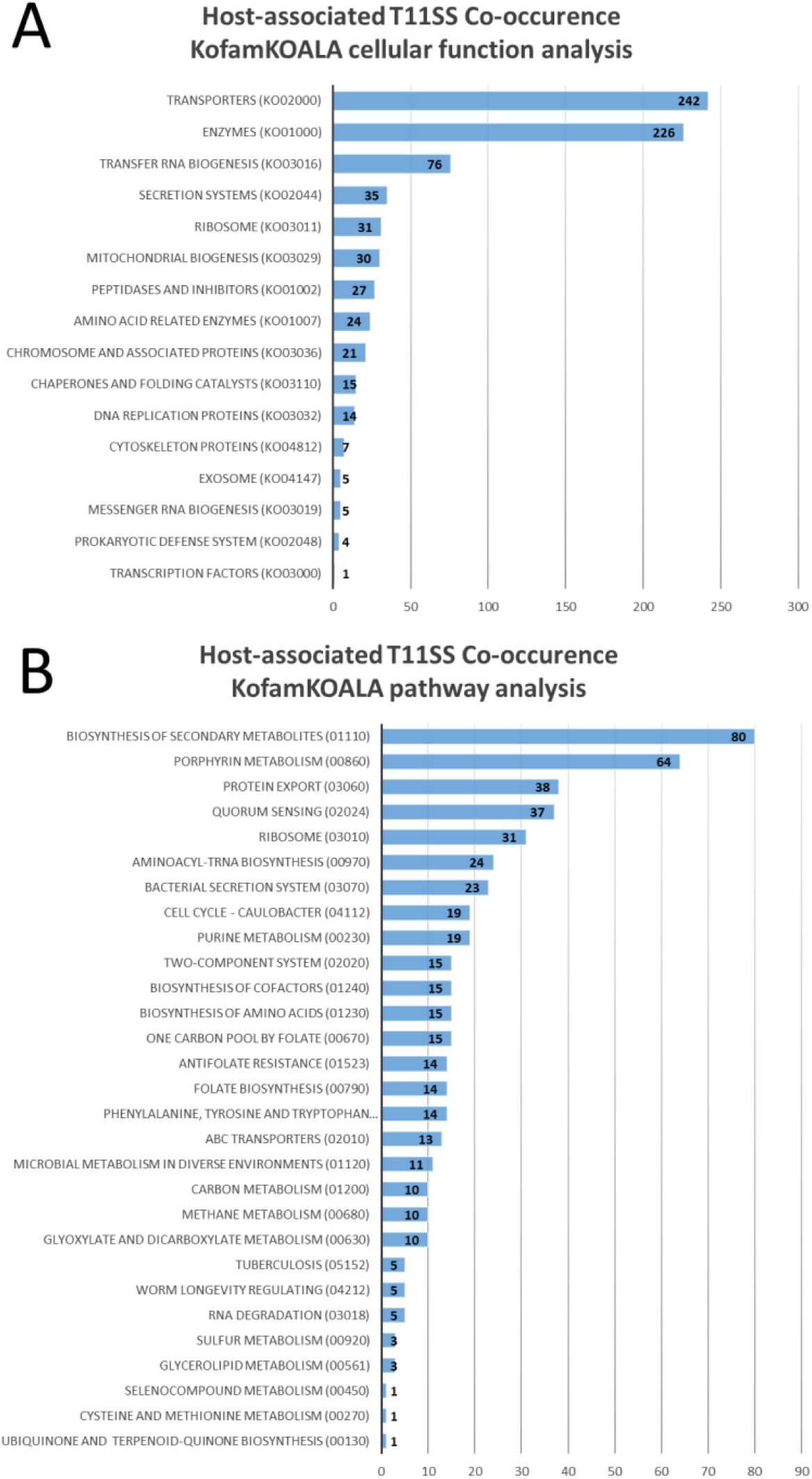
**Host-associated T11SS genome neighborhoods reveal conserved association with iron/heme uptake, export, and one-carbon metabolism**. KofamKOALA uses hidden Markov models to assign functions to query sequences and reveal shared pathways. Cellular functions **(A)** were estimated using BRITE hierarchies and, where possible, assigned to known pathways **(B)** to detect commonalities.

To further functionally annotate co-occurrences, we used BLAST alignment, rather than hidden Markov searching, to establish sequence similarity using BlastKOALA, which compares query sequences to the non-redundant KEGG dataset [25]. The software found matches for 1387 of the 3405 significantly co-occurring genes. Categorization into cellular functions (BRITE hierarchies) again revealed the largest categories to be enzymes, transporters, and tRNA biogenesis (Fig. S3A) (Supplemental file 2, Cluster1BlastBrite tab). BlastKOALA identified more co-occurring amino acid importers (arginine, lysine, tyrosine, tryptophan, and serine transporters) in the dataset than KofamKOALA. Of the 1387 identified genes, 260 had known pathways. This analysis identified similar pathways to those identified by KofamKOALA, including secondary metabolite biosynthesis, protein export, porphyrin metabolism, and carbohydrate biosynthesis (Fig. S3B) (Supplemental file 2, Cluster1BlastPath tab).

### Predicted structure analysis of putative cluster 1 T11SS cargo reveals variable N-termini and shared C-terminal domains

To bioinformatically identify potential cluster 1 T11SS-dependent cargo we searched co-occurring protein sequences for homology to domains of experimentally demonstrated cargo (NilC, TbpB, LbpB, fHbp, HrpC, CrpC, Hpl, and HphA) [4, 7, 12]. The 3405 sequences significantly co-occurring with cluster 1 were first consolidated using UniProt ID mapper into 666 UniRef50 clusters with 50% or greater identity at an amino acid level (Supplemental file 1, Cluster1Uniref tab) [26]. The reference sequences of these UniRef50 were next searched using BLASTp for sequence similarity with N-terminal domains and C-terminal β-barrel domains from experimentally verified T11SS cargo proteins: NilC, TbpB, LbpB, HrpC, CrpC, Hpl, and Factor H binding protein, with a low stringency E-value cutoff of 0.05 [4, 7, 8]. No matches were found for NilC or the N-terminal ligand binding domain of fHbp, but all other queries matched at least one cluster. This analysis revealed 141 clusters of potential T11SS-dependent cargo proteins representing 2656 ORFs across 1048 species/taxa (Supplemental file 1, Cluster1PutativeCargo tab). Each sequence was also annotated with SignalP 6 [37, 38] which predicted 81 Sec secreted lipoproteins, 33 Sec secreted soluble proteins, and 27 proteins with no predicted signal peptide (3 of which were annotated as fragments). For our purposes, we defined a lipoprotein as any predicted to be membrane anchored via tri-acylation of a C-terminal cysteine. Soluble proteins were defined as those predicted to have a Sec (or Tat) signal peptide and that lacked predicted transmembrane features. Overall, the fact that the majority of identified T11SS cargo candidates are predicted to have a signal peptide is consistent with published evidence that T11SS transport Sec dependent lipoproteins and soluble proteins [4, 7].

We utilized multiple sequence alignment to separate the predicted cargo protein clusters into distinct, annotated architectures (secondary structures in Fig 3, tertiary structures in Fig 4, Supplemental file 1, Cluster1PutativeCargo tab). This analysis identified two distinctive groups of Hemophilin-like proteins based on homology to known hemophilins (Hpl, HrpC, HsmA), the predominantly soluble group 1 (96 lipoproteins, 912 soluble) and the predominantly lipidated group 2 (65 lipoproteins, 7 soluble), a group of TbpB-like proteins (638 lipoproteins, 7 soluble), a group of proteins annotated as fHbp-like proteins by virtue of their Lipoprotein C domain (54 lipoproteins, 285 soluble), and a group of lactoferrin-binding protein-like proteins (105 lipoproteins).

**Figure 3.**
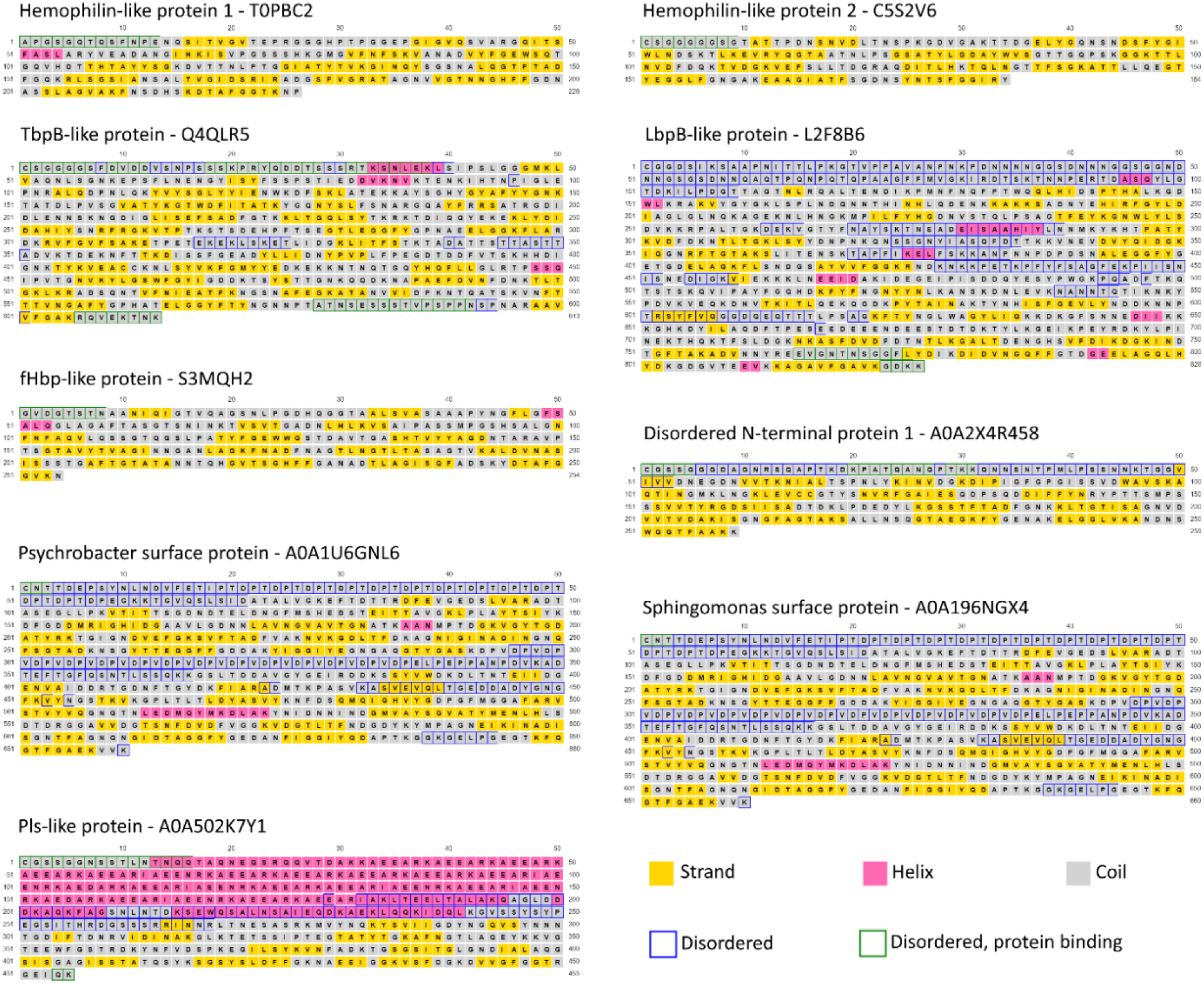
**Secondary structure of predicted T11SS-dependent cargo reveals diverse and novel gene architectures and effector domains**. PsiPred 4 and DisoPred 3 use amino acid sequence to annotate secondary structures highlighting a diversity of N-terminal domains with proline rich repeats, intrinsically disordered regions, α-helical repeats, and ligand binding handles. For each of the detected cargo architectures the representative sequence from the largest UniRef50 cluster was trimmed of it signal peptide to reflect a mature protein.

**Figure 4.**
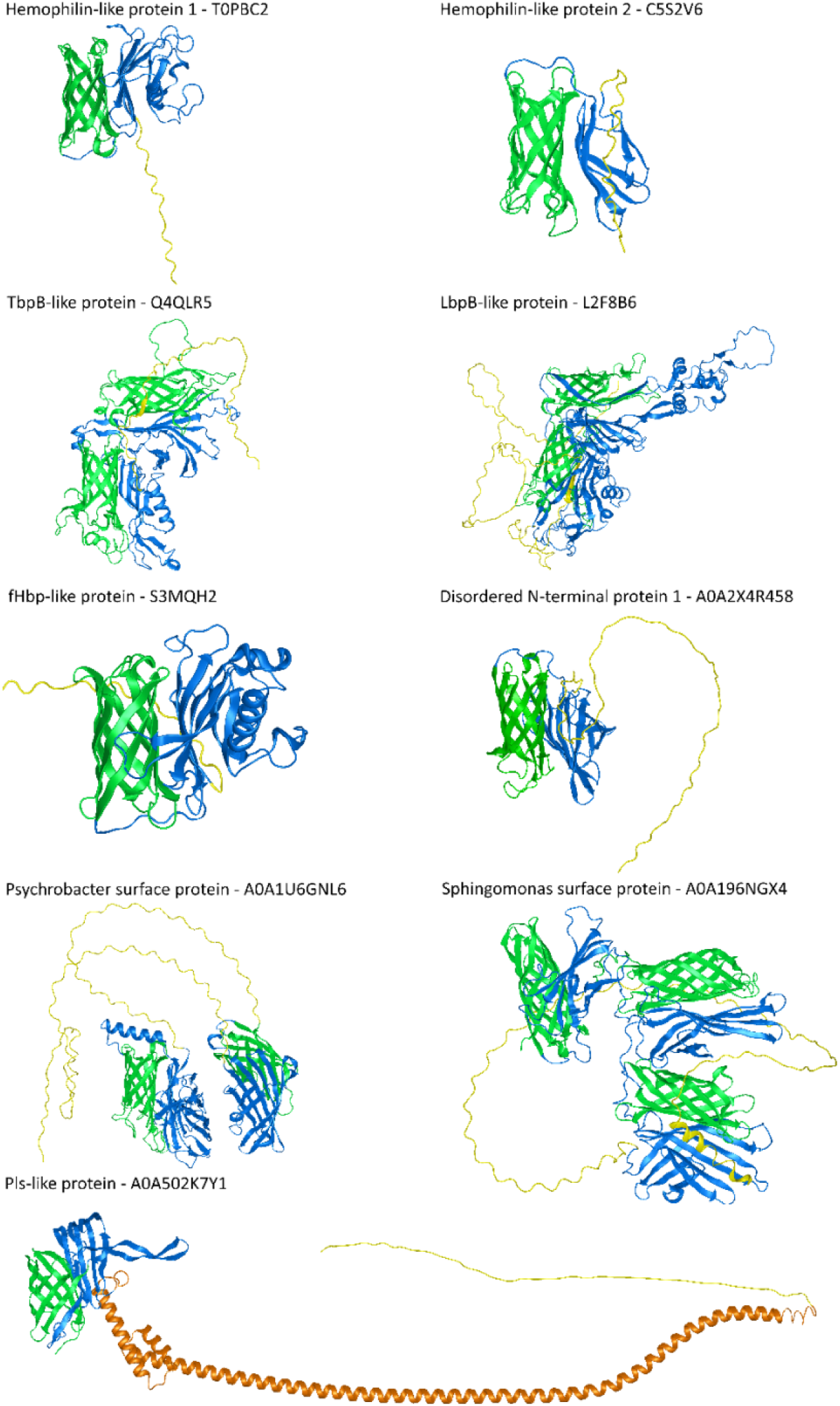
**AlphaFold2 annotation of predicted T11SS-dependent cargo reveals conservation of C-terminal or centrally located β-barrel domains**. For each of the detected cargo architectures the representative sequence from the largest UniRef50 cluster was used to predict a representative structure. Green residues indicate C-terminal β-barrel domains. Blue residues indicate putative ligand binding handle domains. Orange residues indicate glycosylation motif repeats within the Pls-like protein architecture. Yellow residues indicate regions predicted to be disordered.

Several categories with no characterized representatives also emerged. Two lipoprotein architectures with an elongated, disordered N-terminal region occurred: disordered N-terminus 1 (196 lipoproteins, 8 soluble) and 2 (1 lipoprotein only). Another group was named Pls-like proteins (173 lipoproteins, 20 soluble) because of homology to the repeat rich (EEARKA motif) α-helix from Pls surface glycoproteins in *Staphylococcus aureus* . Two phylogenetically-restricted architectures were observed with multiple TbpBBD domains, *Sphingomonas* surface proteins, which have an N-terminal sequence of proline-threonine repeats (18 lipoproteins, 12 soluble) and *Psychrobacter* surface proteins which have an N-terminal sequence of ∼12 proline-threonine-aspartic acid repeats (PTD motif) (55 lipoproteins, 4 soluble). Proline-threonine repeats occur in *Xanthomonas campestris* endoglucanase, where they provide elasticity and facilitate enzyme positioning around its substrate [39]. The PTD motif is found in a family of *Gracilibacteria* (BD1-5) surface proteins and in the eukaryotic microbe *Plasmodium falciparum* and are hypothesized to act as adhesins facilitating interactions with hosts [40, 41].

Using PsiPred 4.0 [27] and AlphaFold2 [21] we predicted the secondary and tertiary structures of representative proteins from the largest UniRef50 cluster from each architecture with more than one representative (Fig. 3 and Fig. 4). All of the architectures were predicted to have at least one domain, generally at the C-terminus, composed of β-strands linked by disordered loops that form a C-terminal β-barrel domain similar to the TbpBBD domain or the Lipoprotein C domain. The N-terminal domains of most cargo had β-strands predicted to form handle domains like those seen in hemophilin or TbpB [4, 7]. When present, the disordered regions and long α-helical repeat regions preceded this handle domain.

When examining secondary and tertiary structures in parallel we noticed that the predicted cargo form three distinctive structural regions: a common C-terminal 8-stranded β-barrel domain, a common central region formed by β-strands, which we have termed a “pedestal” region based on its positioning as a platform for a variable N-terminal domain “effector” domain, predicted to serve the functional role specific to each cargo protein type.

Using solved crystal structures where available [6, 10, 42], and Alphafold2 or Alphafold3 models for representative cargo proteins [21], we divided the predicted cargo structures into these three regions and categorized them by recurrent patterns (Fig. 5). The predicted central pedestal-domains fell into six visibly distinguishable categories; ; 4-stranded groove (Fig. 5A), wing (Fig. 5B-D), and sheet (Fig. 5E, F), 5-stranded sheet (Fig. 5G-I, and 6-stranded/extended sheets (Fig. 5J, K). None of the examined pedestal regions featured complex, variable loops, suggesting they may play a structural role. In contrast, the C-terminal 8-stranded β-barrel core structures are embellished within each protein by diverse inter-strand loops including disordered regions, extended β-strands, and α-helices that could be playing functional roles, as do the loops of the N-lobe TbpBBD domain of TbpB [43].

**Figure 5.**
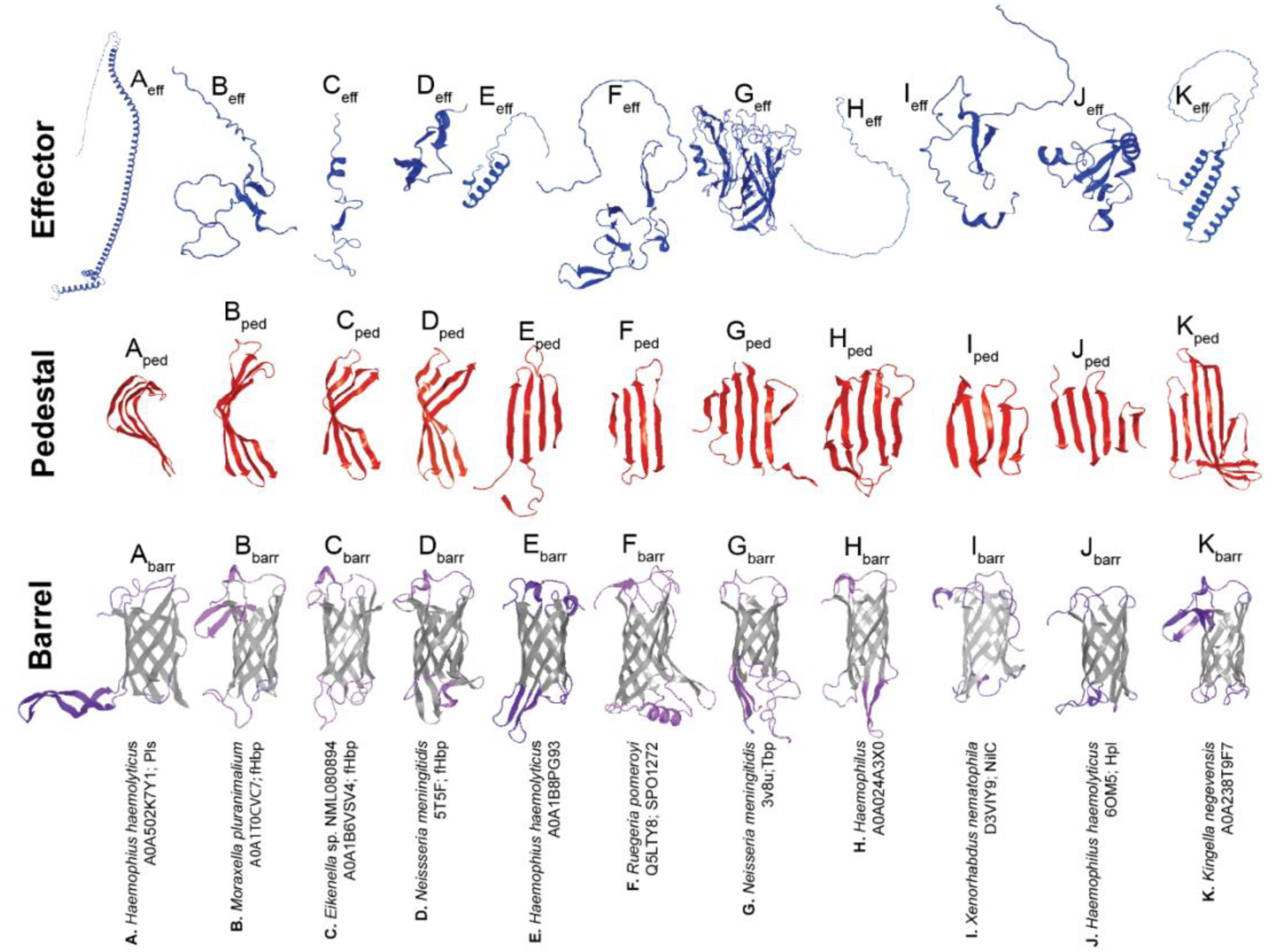
AlphaFold2 annotation of predicted T11SS-dependent cargo reveals three distinct domains ubiquitous to cargo. Solved **(D, G, J)** or predicted **(A, B, C, E, F, H, I, K)** structures from representative UniRefF50 cluster sequences of predicted **(A, B, D, E, F, H, K)** or known cargo **(D, G, I, J)**. Each structure was separated into three sections representing common structural domains: N-terminal effector (blue), pedestal (red), and C-terminal β-barrel (gray barrel residues and purple loop residues). N-terminal effector domains varied in their predicted structural motifs and included disordered **(B, C, D, H, I)**, α-helical **(A, E, K)**, β-strand **(D, F, J)**, and barrel **(G)**. Pedestal domains comprised 4-stranded groove **(A)**, 4-stranded “wing” **(B-D),** 4-stranded sheet **(E, F)**, 5-stranded sheet **(G-I)**, or 6-stranded/extended β-sheet **(J, K)**. Some of these β-strands are donated by primary sequences from either the N-terminus or the C-terminus of the protein. Barrel domains exhibited variation in the presence of extended loops, helices, or strands in predicted surface- or periplasmic-exposed regions, as indicated by purple coloration.

The N-terminal effector domains displayed a variety of structural motifs including disordered loops (Fig. 5 B-D,H, I), α-helices (Fig. 5A, E, K), β-strands (Fig. 5D,F,J), and β-barrels (Fig. 5G). The examined effector regions ranged in size, with some barely extending beyond the pedestal domain and others including duplications of the 8-stranded β-barrel domain. The latter is typified by the solved structure of TbpB, which comprises two lobes, each containing the hemophilin handle-β-barrel pair architecture (Fig. 5G). The TbpB N-lobe pair comprises the effector region, while the C-lobe handle comprises the pedestal and the C-lobe barrel is the putative T11SS targeting domain [42, 43]. This domain duplication and functional specialization may also explain the architecture of *Sphingomonas* and *Psychrobacter* surface proteins that feature 2-3 copies of the handle-barrel pair (Fig. 4). However, the long, disordered loops present in those proteins likely generate a distinctive structural architecture relative to TbpB.

### Demonstrating T11SS-dependent secretion of Plasmin sensitive surface protein

To assess the accuracy of our bioinformatic predictions of T11SS cargo, we experimentally tested levels of secretion in *E. coli* of a representative novel cargo protein with or without its predicted cognate T11SS protein. Several of our predicted cargo possess a large α-helix repeat region absent from the known T11SS cargo. We chose to test a representative of this group, the 525 amino acid <u>P</u>lasmin <u>s</u>ensitive <u>p</u>rotein homolog (Pls) (STO63610.1) and its genomically associated putative type <u>e</u>leven <u>P</u>ls <u>s</u>ecretor (TepS) (STO63613.1) from *H. parahaemolyticus* strain NCTC10794. The *H. parahaemolyticus* Pls α-helix repeat region shares between 49 and 62% identity with the repeat regions of surface glycoprotein Pls from *Staphylococcus aureus* (WP_256928426, WP_256927786, WP_258808513, WP_256933640, and WP_257570445) that contributes to virulence by stimulating biofilm formation [44]. *Staphylococcus* and *H. parahaemolyticus* Pls differ in that homologs of the former are larger and lack the C-terminal β-barrel domain.

To experimentally test for TepS-mediated secretion of *H. parahaemolyticus* Pls, we used addition of a FLAG-tag epitope to allow Pls detection and expressed this protein with or without TepS using pETDuet-1 based plasmids in *E. coli* BL21 DE3 C43. Western blot analysis of lysed cells revealed that Pls was effectively expressed with or without TepS co-expression, though slightly more was present in/on cells expressing TepS, suggesting that surface exposure may increase the amount of protein a cell can contain, or that TepS protects the protein from degradation (Fig. 6A and S4AB). Flow cytometry was used to detect surface exposure of membrane anchored lipoproteins. No Pls was detected on the surface of cells induced to express Pls alone, however ∼40% of cells had detectable surface Pls when expressing both Pls and TepS (Fig. 6BC). Since Pls homologs can be cleaved spontaneously, or by proteases as in the case of *S. aureus* Pls [44, 45] we used western blot immunostaining on spent culture media (separated into soluble and vesicle fractions) to determine if Pls was being liberated from cell surfaces (Figure 6D and S4CD). Immuno-dot-blots were performed on cell lysates after collecting supernatants to ensure that comparable amounts of Pls were being produced (Fig. 6D and S4E) [44, 45]Pls was present in the supernatant, but only when co-expressed with the T11SS protein TepS, suggesting that *Haemophilus* Pls can be cleaved from the cell surface spontaneously. [44, 45]Some Pls was located in extracellular vesicles independent of TepS expression, but this amount was higher in the presence of TepS (Fig. 6 and Fig. S4). In Western immunoblots, FLAG-tagged Pls is visible as a single clear band in individual replicates, but varied in apparent size between biological replicates from 82.3 to 121.3 kD with an average of 103kD. This is almost twice the value predicted from sequence alone (56.8kD). This large and variable size may suggest that Pls is either being significantly slowed by glycan moieties as it travels through polyacrylamide, its passage is being affected by its extremely high concentration of charged amino acids, or that it exists as a dimer when expressed in *E. coli*.

**Figure 6.**
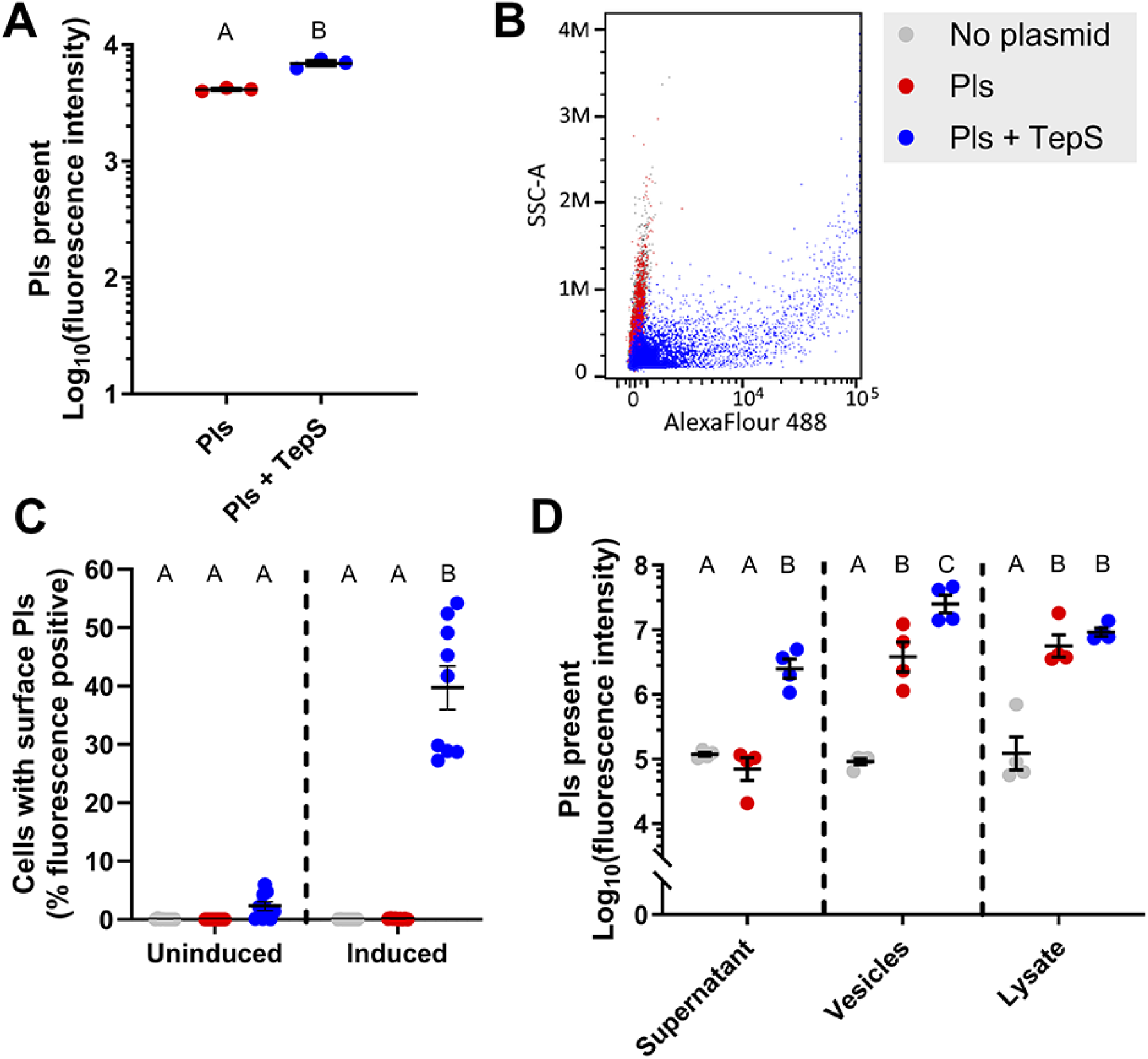
Localization of FLAG tagged Pls in the absence and presence of its cognate T11SS secretor. *E. coli* cells expressing Pls-FLAG alone or Pls-FLAG alongside its cognate T11SS (TepS) were probed using Western blotting (lysates from flow cytometry samples, supernatants, and extracellular vesicles), flow cytometry (cell surface exposure), and immuno-dot blots (lysates from supernatant collections) to see how expression of the T11SS protein impacted protein localization. Strong expression of Pls was detected in lysates from both strains **(A and D)**. A representative flow cytometry replicate **(B)** shows that surface exposed Pls was only detectable on cells co-expressing TepS **(C)**. For each biological replicate 100,000 events were detected via flow cytometry. Supernatant secretion of Pls was also only detectable in the presence of TepS **(D)**. The extracellular vesicle fraction collected via ultracentrifugation showed some Pls localization in the absence of TepS, however that localization was significantly increased by TepS. One-way ANOVA with Tukey’s HSD statistical test used to compare treatments in all experiments, letters above treatments indicate significance groups at p value 0.05. Error bars indicate standard error of the mean.

### Bioinformatic analysis of cluster 3 T11SS proteins from marine environments reveals distinct co-occurrence patterns

To test the robustness of our analysis we then utilized our co-occurrence technique to explore cluster 3 T11SS proteins, predominantly containing sequences from marine microbes. Cluster 3 was chosen for in depth analysis because it is composed entirely of sequences from organisms that occupy aquatic/marine environments, for which no T11SS have been described in the literature and which may encode novel cargo. This cluster almost exclusively contains sequences from the families *Roseobacteriaceae* and *Rhodobacteraceae* [46], including both pelagic microbes and symbionts of algae (*Silicimonas algicola*) [47], mollusks (*Aliiroseovarius crassostreae*) [48], tunicates (*Ascidiaceihabitans donghaensis*) [49], echinoderms (*Sulfitobacter delicatus*) [50], and corals (*Roseivivax isoporae*) [51]. All genes from cluster 3 (145 sequences) were used as queries in the RODEO genome neighborhood network function, using the same parameters as the cluster 1 analysis, resulting in 203 co-occurring domains with frequencies between 2 and 102 (Supplemental file 1, Cluster3Co tab). Domains which co-occurred fewer than 5 times were excluded from analysis (122 or ∼60.0%) (Fig. S1C-S1D). Co-occurring domains were compared to another random subset of the non-specific control co-occurrence dataset to assign FDR values. All domains with an FDR greater than 0.1 were excluded from analysis, resulting in 42 significantly co-occurring domains (Supplemental file 1, Cluster3FDR_Threshold tab). The most common co-occurring domains were additional DUF560/T11SS domains (102/145 loci), DUF1194 (102/145 loci), glyoxalase domains (89/145 loci), and thymidylate synthase/thymidylate synthase complementing protein (77/145 loci). The TbpBBD domain passed the FDR threshold, however relative to the analysis of cluster 1 it was exceptionally rare (8/145 loci). All genes that co-occurred with a marine cluster 3 T11SS OMP and contained any of the significant co-occurring domains were extracted, resulting in 860 significant co-occurring genes which we submitted to KofamKOALA and BlastKOALA for functional analysis [24, 25].

KofamKOALA found matches for 595 of the 860 significant co-occurring genes (Supplemental file 2, Cluster3KofamBrite tab). When matched to cellular functions (BRITE hierarchies) the most common categories were enzymes (lactoylglutathione lyase, thymidylate synthase complementing proteins, aspartate aminotransferase, etc.), prokaryotic defense systems (antitoxins CptB and HigA-1, topoisomerase IV B), and transcription factors (Cell division repressor DicA, cold shock protein) (Fig. 7A). Of the 595 matched sequences, 270 had known pathway association, the most common pathways being biosynthesis of one carbon pool by folate (dihydrofolate reductase, thymidylate synthase), pyrimidine/nucleotide metabolism (thymidylate synthase), pyruvate metabolism (lactoylglutathione lyase), and biosynthesis of amino acids (aspartate aminotransferase) (Supplemental file 2, Cluster3KofamPath tab). Unlike cluster 1, transposases were not among the significantly co-occurring genes of cluster 3.

**Figure 7.**
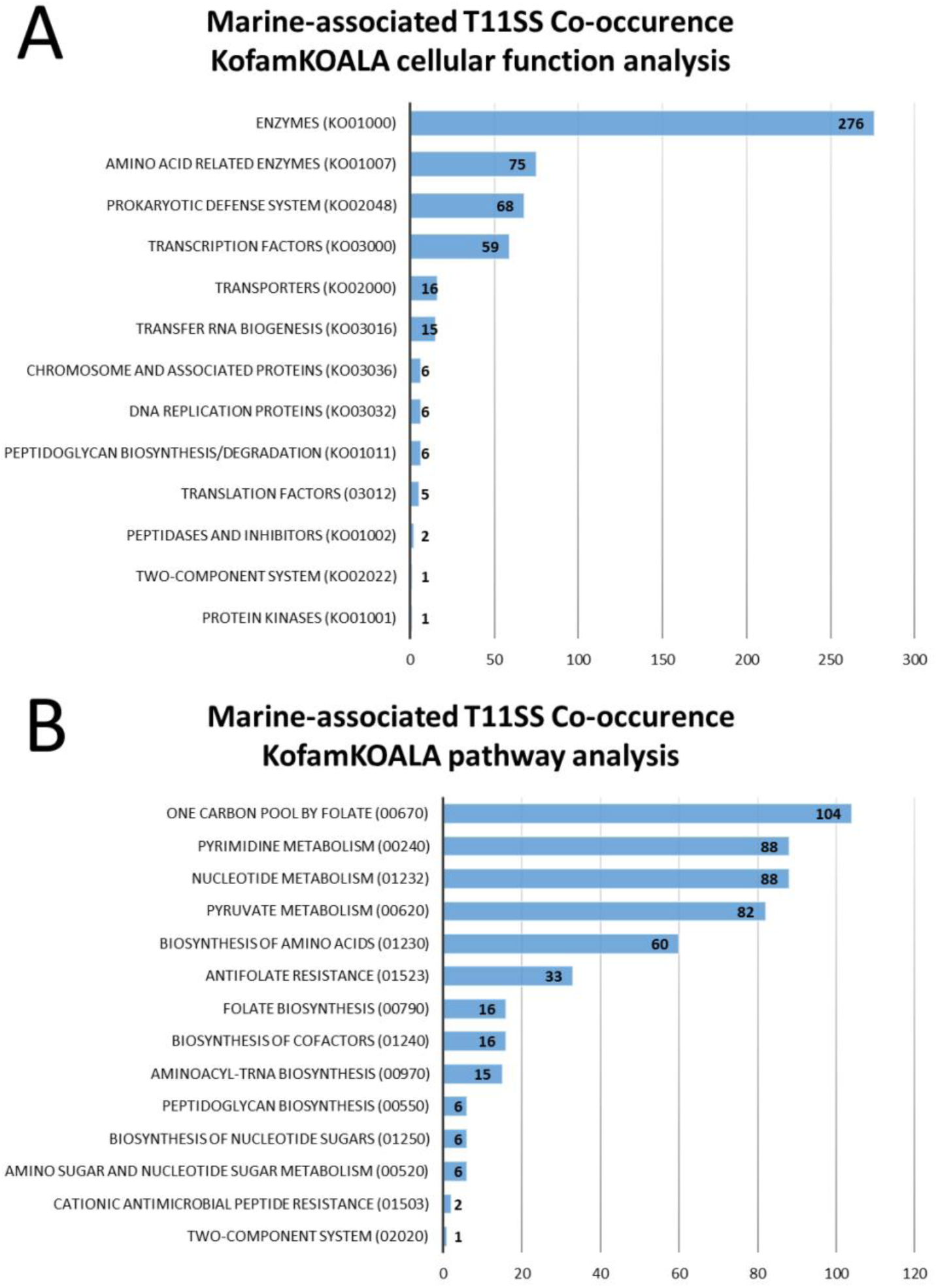
**Marine T11SS genome neighborhoods reveal association with one-carbon metabolism, nucleotide metabolism, and the glyoxalase-detoxification pathway.** KofamKOALA uses hidden Markov models to assign functions to query sequences and reveal shared pathways. Cellular functions **(A)** were estimated using BRITE hierarchies and assigned, where possible, to known pathways**(B)**.

**Figure 8.**
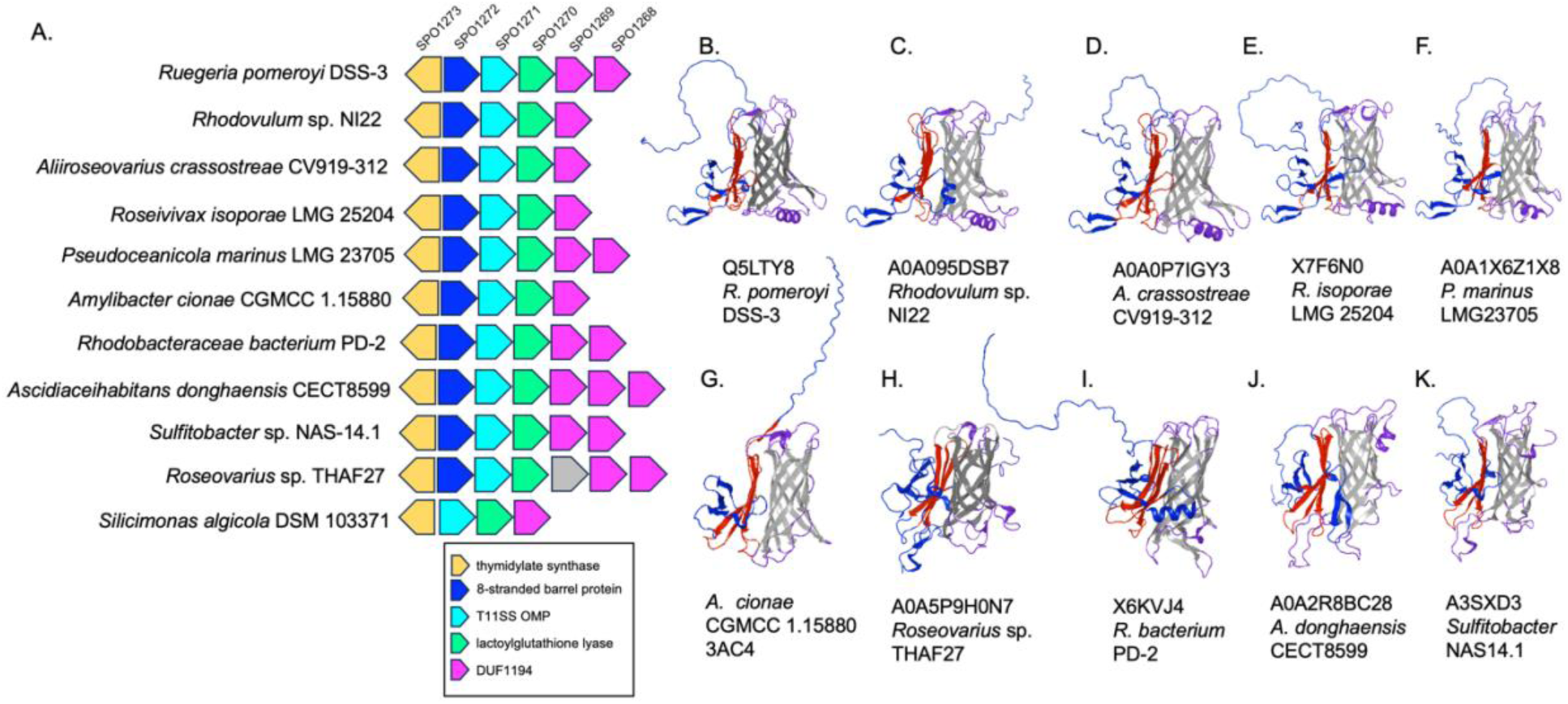
A conserved marine cluster 3, T11SS OMP locus encodes predicted cargo proteins. A schematic representation of select examples of a T11SS OMP-encoding, syntenic locus found with genomes of indicated strains of *Rhodobacteraceae* and *Roseobacteriaceae* **(A)**. Each gene is represented by a block arrow (not to scale), and the *Ruegeria pomeroyi* locus tags of each homolog are indicated at the top. In all 11 genomes examined, the syntenic locus included genes predicted to encode a T11SS OMP (turquoise arrows) as well as the top cluster 3 co-occurring functions: flavin-dependent thymidylate synthase (gold arrows), lactoylglutathione lyase (neon green arrows), and one or more DUF1194/von Willebrand factor proteins (pink arrows). In all but one, the locus includes a putative T11SS cargo protein (light blue arrows), identified based on the presence of a predicted C-terminal 8-stranded β-barrel domain. Predicted AlphaFold 2.0 or AlphaFold 3.0 predicted structures of putative cargo proteins encoded within these loci is shown in **(B-K)**. Color coding is used to indicate the putative effector (dark blue), pedestal (red), barrel (gray), and surface or periplasmic loops (purple) of each protein. The top five appear to have similar structural motifs, including an α-helix in one of the β-barrel loops, antiparallel β-strands in the N-terminal domain (blue), and a 4-stranded sheet pedestal domain (red). The remaining five structures show the overall domains of N-terminal (blue), pedestal (red) and β-barrel (gray), but the N-terminal domains are predicted to have different structural motifs relative to the others examined.

However, aminoacyl-tRNA synthesis functions, which can be associated themselves with mobile genetic elements [36, 52], do appear on the co-occurrence list, including alanyl-tRNA, prolyl-tRNA, and threonyl-tRNA synthases (Fig. 7B). Parallel analysis with BlastKOALA identified 514 of the 860 significant co-occurring genes, of which 267 had known pathway association (Supplemental file 2, Cluster3BlastBrite and Path tabs). BlastKOALA did not identify any additional cellular functions or pathways not already revealed by KofamKOALA (Fig. S5).

### Bioinformatic investigation of potential cluster 3 T11SS cargo

Initial attempts to identify new cargo proteins within the co-occurrence datasets from cluster 3 used the same methods used for cluster 1. UniRef50 clustering with ≥50% identity resulted in 218 homologous groups (Supplemental file 1, Cluster3Uniref tab) that were assessed with BLASTp for sequence level homology to experimentally characterized T11SS cargo domains. Only a few sets of homologs were identified, even when including all TbpBBD-domain-containing UniRef50 clusters. Homologs of six UniRef50 clusters, specific to the *Rhodobacteraceae*/*Roseobacteriaceae* families, resembled the Hemophilin-like 1 architecture, and homologs from another UniRef50 resembled the disordered N-terminus 1 architecture noted previously (see Figs. 3 and 4). Collectively these 7 UniRef50 clusters encompass a total of only 31 ORFs across 27 species/taxa (Supplemental file 1, Cluster3PutativeCargo tab). The ratio of predicted cargo per T11SS OMP was far lower in cluster 3 (0.21 Cargo/input OMP) than it was in cluster 1 (2.39 Cargo/input OMP). This dearth of co-occurring putative cargo proteins suggests that marine cluster 3 T11SS OMPs seldom co-occur with their cognate cargo proteins, utilize cargo proteins with below-threshold sequence similarity to those of cluster 1 T11SS, or have evolved distinct functions and no longer have partner cargo proteins.

To further examine this potential transition in function, we comparatively annotated a syntenic T11SS OMP-encoding locus in 11 representative *Rhodobacteraceae*/*Roseobacteriaceae* strains (Supplemental file 3, Clus3SyntenicPairs tab). Each locus encodes all four of the top co-occurring domains from our co-occurrence analysis, including flavin-dependent thymidylate synthase, lactoylglutathione lyase, and one or more DUF1194 (Fig. 8A and Supplemental file 3, NeighborhoodAlignment tab). All but one (*Pseudooceanicola marinus* LMG23705) of the loci also encodes a protein that is predicted to be a T11SS cargo lipoprotein (282-375 aa) based on the presence of an N-terminal SPII type signal sequence and a C-terminal 8-stranded β-barrel domain (Fig. 5F; Fig. 8B-K) [21]. The presence of both cluster 1 and cluster 3 co-occurring domains indicates that this conserved locus is a hybrid of the two. It is tempting to speculate that this locus represents a transitional state in the divergence of these systems, with at least one analyzed genome appearing to have lost the cluster 1 co-occurring, 8-stranded β-barrel domain.

We next examined the cluster 3 marine T11SS co-occurrence dataset to consider the possibility of alternative putative cargo types. In contrast to cluster 1 T11SS, co-occurrence with TbpBBD/lipoprotein C domains, TonB-dependent receptors, and TonB domains are rare within the marine cluster 3 T11SS loci. Instead, the DUF1194 domain (PF06707) was highly prevalent (102/145 queries) (Supplemental file 4). Of the 102 DUF1194-containing sequences found in the cluster 3 co-occurrence list, 7 are predicted to be lipoproteins, 76 are predicted to be soluble Sec-secreted proteins, and 19 have no detectable signal peptide according to SignalP 6 [37, 38]. Many cluster 3 T11SS loci encoded 2 or 3 distinct DUF1194 genes in close proximity (e.g., Fig. 8), predicted to encode both lipidated and non-lipidated proteins (e.g., algal symbiont *Rhodobacteraceae* PD-2 ETA49263.2 and ETA49262.2, respectively) (Supplemental file 4). The predicted RoseTTAFold structure [53] of the DUF1194 domain is a mixed β-strand/α-helix (Fig. 8). According to the Pfam database, the DUF1194 domain occurs in 31 distinct protein architectures with other domains, including C-terminal autotransporter domain (PF03797) predicted to function in T5SS secretion, the CARDB domain (PF07705), which adopts a 7 β-strand structure, DUF11 (PF01345), predicted to have an 8-stranded β-barrel structure, and PF18911, the known crystal structures of which also adopt a β-barrel (e.g., 1wgo and 2y72).

Of the co-occurring DUF1194 sequences, 6 are annotated as having homology to <u>v</u>on <u>W</u>illebrand factor type <u>A</u> domain or vWA (PF00092). This homology is reflected in the structural similarity between these two domains, including the presence of a Rossman fold, which may hint at similar molecular roles. vWA protein architectures can include divalent-cation-binding <u>m</u>etal ion-<u>d</u>ependent <u>a</u>dhesion <u>s</u>ites (MIDAS), with a characteristic DxSxS…T…D motif [54, 55].

Almost all the DUF1194 proteins that co-occurred with DUF560 proteins contained at least one MIDAS motif (98/102). To examine potential structural features of cluster 3 T11SS co-occurring DUF1194-containing homologs, we focused on three adjacently encoded DUF1194 genes from *Ascidiaceihabitans donghaensis* (SPH20589/SPH20588/SPH20587) for structural analysis. PsiPred 4.0 predicted that these proteins have at least 5 predominantly hydrophobic β-strands, all separated by α-helix regions in a structure that also appears to be a variation on the classical α/β Rossman fold (Fig. 9A). RoseTTAFold predictions of the tertiary structure of these DUF1194 proteins reveal globular proteins wherein 6 hydrophobic β-strands form a β-sheet at the core of the protein, which is protected by three α-helices per side. All three representatives had a MIDAS motif starting 10-11 residues after the predicted signal peptide. For each protein, the last 11-13 C-terminal residues appeared to be disordered, and their positions could not be estimated with a high degree of confidence (Fig. 9B).

**Figure 9.**
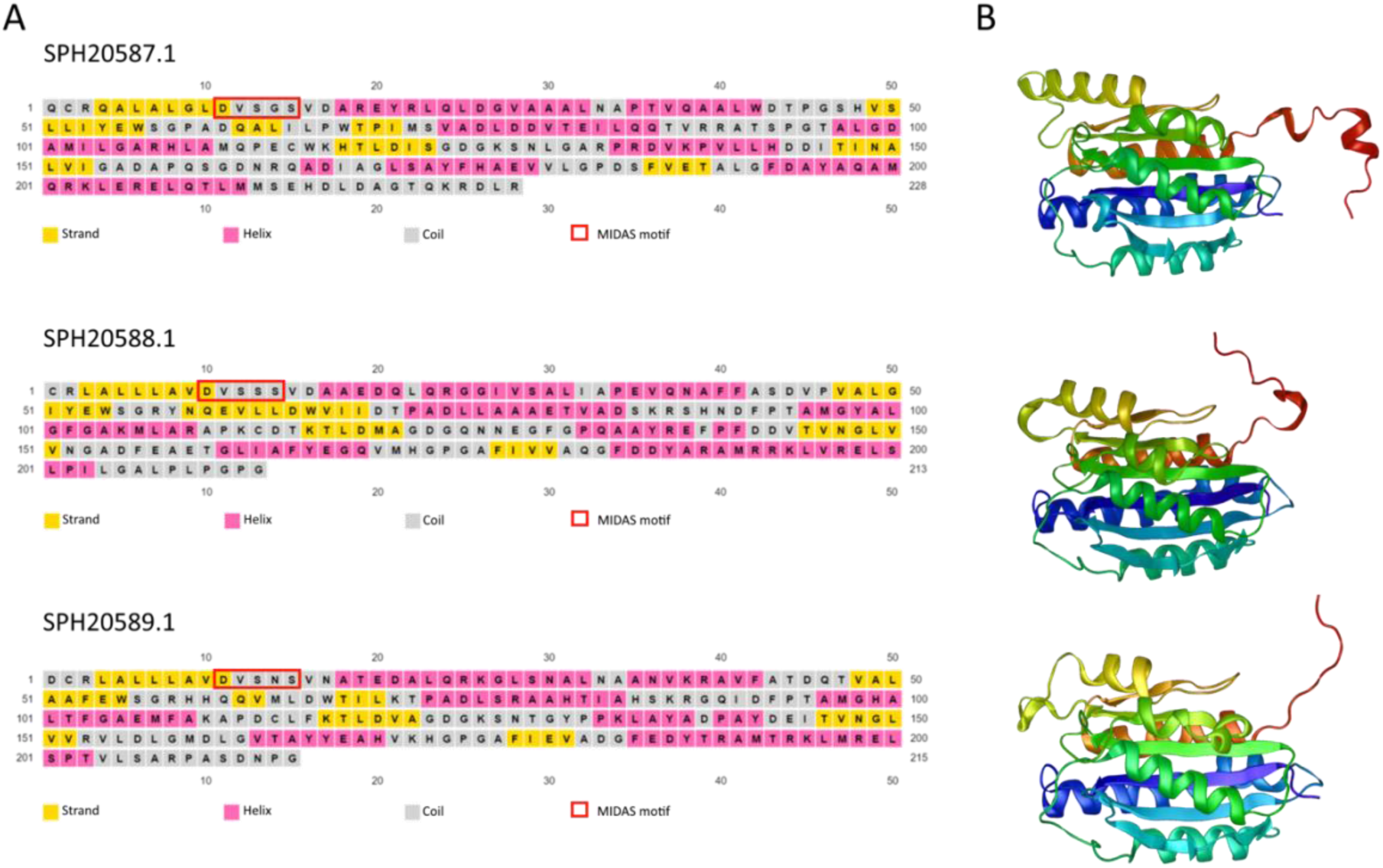
**Representative structures of DUF1194 proteins located within a T11SS genomic neighborhood in *Ascidiaceihabitans donghaensis***. These three DUF1194 homologs are encoded adjacent to a T11SS OMP-encoding gene within a microbe isolated from a tunicate host. These proteins appear to be paralogs of each other based on sequence similarity. Signal peptides were predicted with SignalP 6 and trimmed from the sequences prior to submission to PsiPred **(A)** and RoseTTAFold **(B).**

## Discussion

The controlled bioinformatics analyses we report here reinforce the idea that cluster 1 T11SS-dependent cargo have roles in metal uptake, single carbon metabolism, and nutrient provision. Further, our data extend the experimentally verified T11SS cargo to include *Haemophilus parahaemolyticus* Pls and lend credence to the concept that T11SS cargo can be bioinformatically predicted. The structural prediction that this cargo protein is a fusion between a functional Pls domain, typical of Gram-positive bacteria, and a C-terminal 8-stranded β-barrel domain lends further evidence of the function of the barrel domain in directing cargo to the T11SS, and for the function of T11SS in facilitating transport across an outer membrane. However, our analysis of the more distantly related cluster 3 T11SS OMPs revealed that known barrel domains (TbpBBD and lipoprotein C) are rare among the significantly co-occurring genes identified, raising the intriguing possibility that novel structural domains may exist that mediate targeting to cluster 3 T11SS.

Our bioinformatic approach relied upon gene co-occurrence analysis, which uses genomic proximity as a proxy for coordinate regulation and potential interactions, since genes within a common functional pathway often cluster together within a genome. While this is not a foolproof method, the use of large genomic datasets increases the power of co-occurrence analysis to identify potential co-interacting partners [56]. One of the methodological refinements we developed was the use of *in silico* controls for genome neighborhood co-occurrence to reduce false positives and noise in the output datasets of putative T11SS co-occurring protein families. For instance, activities associated with protein translation and RNA metabolism, which are essential cellular processes expected to be present in every cell, are not always useful co-occurrence signatures for elucidating T11SS function and components. Such general functions are not indicative of the specific functions of T11SS relative to any other OMP and can be considered a form of false negative, Type II error; such co-occurrences are not biologically relevant, but are not being excluded from the analysis because they do co-occur frequently with the query gene. However, removal of such co-occurrences can also lead to false positive, Type I error, because they can also result in removal of some biologically relevant co-occurrences. For example, there could be protein families that are important functional partners of the query protein, but that are removed from the dataset if they were important for both query and control proteins. This type of error occurred in our analysis. For example, TonB-dependent receptors, with which many T11SS-dependent cargo interact, were excluded from the cluster 1 T11SS analyses since these domains associate with other outer membrane proteins unrelated to T11SS. In a sense, this control narrows the focus from total co-occurrence to unique co-occurrence, and as such it is best suited towards applications in which the researcher seeks to differentiate protein families. Additionally, the technique can be further adapted and refined, depending on application. For instance, multiple non-specific gene neighborhood controls could be generated by randomly subsampling the list of proteins that are biophysically similar to the query proteins, allowing for increased stringency and measures of variance.

The cluster 1 T11SS significantly co-occurring genes identified here are consistent with previous observations [4, 7]. Iron uptake from the environment is a common function among experimentally verified T11SS-dependent cargo proteins and this theme was extended within the cluster 1 T11SS co-occurrence dataset, which included heme oxygenases and heme transporters that facilitate iron uptake [57]. Similarly, the protein export genes identified as co-occurring with cluster 1 T11SS included TonB and ExbD, which are essential to energize uptake of nutrients from known T11SS-dependent cargo [8, 10, 58]. TolA is essential to bring periplasmic nutrients into the cytoplasm of the cell [59]. Signal peptidase II is essential for the generation of bacterial lipoproteins, which constitute a substantial percentage of T11SS-dependent cargo [60]. Finally, the observed co-occurrence with transposases and tRNA-synthases is consistent with the hypothesis that T11SS are regularly associated with mobile genetic islands and can be horizontally acquired and contribute to fitness within a host environment [36].

Significantly co-occurring pathways were commonly identified for both cluster 1 and cluster 3 T11SS, and represent pathways not previously associated with T11SS. These included folate (vitamin B9)-related pathways, including folate biosynthesis, folate-dependent enzymes, and one carbon metabolism via folate. Folate, like heme, is an enzyme cofactor required for many diverse biological pathways. It is essential for the function of thymidylate synthase complementing protein, also known as flavin-dependent thymidylate synthase, in nucleotide biosynthesis [61]. This protein is the third most common cluster 3 marine T11SS co-occurring gene, suggesting coordinate regulation or inheritance. Since, unlike heme, folate does not incorporate a metallic ion and most bacteria synthesize their folate instead of scavenging it from the environment [62], we do not favor the hypothesis that folate itself is acquired through a T11SS-dependent uptake mechanism. Instead, the close association of folate metabolism and T11SS may reflect the fact that folate biosynthesis, as well as iron-sulfur cluster biosynthesis, and methylotrophic metabolism by formate dehydrogenase (which also significantly co-occurred with T11SS) depend on the metal-bearing enzyme cofactors, cobalamin (vitamin B12) and molybdopterin cofactor (MoCo) [63–66]. Our data suggest that some T11SS OMPs may function in cobalamin or molybdopterin uptake pathways. Regardless, our data clearly demonstrate an association between T11SS and single carbon metabolism and methyltropism.

Cluster 1 T11SS co-occurrences featured TonB energization or heme/iron uptake that were lacking from cluster 3 T11SS co-occurrences. In turn, the most frequent cluster 3 T11SS co-occurrences were DUF1194 and lactoylglutathione lyase (GloA), neither of which appeared in the Cluster 1 T11SS co-occurrence datasets, but both of which have connections to metal ions. The function of DUF1194 is currently unknown but it is predicted to fold into a globular protein with an α/β Rossman fold.

Several of the DUF1194 proteins (6/103) we detected also had homology to a structurally similar α/β Rossman fold domain called von Willebrand factor A (vWA), which is most often associated with serum glycoproteins and large multimeric protein complexes in mammalian systems [67, 68]. In *Myxococcus xanthus* bacteria, the vWA domain-containing surface lipoprotein CglB acts as a surface adhesin essential for gliding motility [69]. Upon binding a ligand, vWA domain-containing adhesins such as CglB can stabilize their bond via a conformational change induced by MIDAS motif allosteric binding of a divalent metal cation, often Mg^2+^[55, 69]. The presence of a MIDAS motif in the vast majority of DUF1194 identified within our dataset suggests that these proteins may also play a role in adhesion or may participate in metal homeostasis in previously unpredicted ways. Additionally, vWA domain containing proteins are involved in microbial conflict sensor systems called MoxR-vWA-centric ternary systems, wherein a surface receptor protein receives an extracellular signal from a symbiotic species and then signals a response using intracellular vWA chaperones and MoxR ATPases [70, 71]. Although we did not detect MoxR homologs in our analysis, this example raises the possibility that one role for T11SS OMPs may be to deliver sensor proteins to the cell surface that relay signals to DUF1194 partners. Other possibilities, include that DUF1194 proteins themselves are a type of T11SS-dependent cargo, or that they contribute to a multimeric protein complex that interacts with T11SS proteins. Lactoylglutathione lyase is an enzyme with a coordinated metal ion (typically nickel) and functions in the regeneration step of the glyoxalase-mediated aldehyde detoxification system [72]. It is possible that the nickel requirement of this enzyme reflects a role for cluster 3 T11SS in facilitating nickel uptake.

Alternatively, this co-occurrence may indicate that cluster 3 T11SS are more generally participating in an aldehyde producing process. Notably, aldehydes like methylglyoxal are exceptionally cytotoxic and can damage DNA and proteins, so the cluster 3 T11SS co-occurrence of nucleotide and amino acid biosynthesis may reflect a more general relationship between T11SS and the aldehyde repair response [72, 73].

In addition to providing insights into functional pathways associated with T11SS, our analysis also allowed us to identify 148 homology groups of putative T11SS-dependent cargo, including those with domain architectures distinct from those found in known cargo. We verified that at least one of these, the <u>pl</u>asmin-<u>s</u>ensitive surface protein (Pls) from the Gram-negative organism *H. parahaemolyticus,* is a bona fide T11SS-dependent cargo protein, demonstrating the efficacy of our bioinformatics approach. Pls represents a novel N-terminal effector domain among T11SS cargo. This domain is homologous to glycosylated Pls proteins found on or spontaneously cleaved from the surfaces of Gram-positive organisms such as *Staphylococcus aureus* and *S. epidermidis* where function in biofilm formation and modulating adhesion to host proteins [44, 45]. Given that the function of Pls requires surface exposure, its presence in a Gram-negative organism is complicated by the existence of the outer membrane. In *H. parahaemolyticus*, this hurdle seems to have been overcome via fusion of the Pls repeat domain to the C-terminal 8-stranded β-barrel domain necessary for cluster 1 T11SS-dependent secretion across the outer membrane [4, 7].

When co-expressed in *E. coli* with its T11SS partner, TepS, Pls predominantly localizes to the supernatant and is not tethered to the cell surface as would be expected of a lipoprotein. Our working model is that some Pls can reach the cell surface in the absence of T11SS, as has been observed for factor H binding protein [74, 75], and that in the presence of the T11SS TepS, Pls is secreted but then cleaved, releasing it into the extracellular milieu. Given that our experiments were conducted in a protease-deficient strain of *E. coli*, it will be exciting in future studies to determine if TepS alters the topology of Pls in order to make it susceptible to proteolysis, or if it is directly responsible for Pls cleavage, which would be a novel function for T11SS. The possibility that Gram-negative Pls are secreted via T11SS and may be released from the cell surface in the process will be an important factor to consider in ongoing efforts to target Pls homologs for vaccine development, such as that being pursued for *Actinobacillus pleuropneumoniae* Pls (YP_001652736.1) [76]. Also, the Pls-like proteins have repeat rich regions and the possible glycosylation of *H. parahaemolyticus* Pls is supported by our observations of its aberrant mobility in SDS-PAGE. It is possible that other T11SS-dependent cargo, including lipoproteins, may also be glycosylated. If so, such glycoproteins tethered to the cell surface would display glycans as surface antigens that may modulate host-microbe recognition and immune evasion [77–79].

In summary, our findings have enhanced our understanding of potential effectors of the novel T11SS and expanded the list of experimentally verified T11SS secreted cargo. We also have expanded the field of genomic co-occurrence analysis by establishing a protocol that can serve as a basis for more controlled and specific experimental designs. Exciting avenues of future research could focus on generating an automated pipeline for controlling co-occurrences analyses, determining if Pls is required for host-colonization by *Haemophilus parahaemolyticus*, investigating if DUF1194 represents a novel T11SS cargo type, and exploring the relationship between one carbon metabolism and T11SS.

## Conflicts of interest

The author(s) declare that there are no conflicts of interest.

## Funding information

This work was funded, in part, by funds awarded to H.G.-B. from the University of Tennessee-Knoxville (David and Sandra White Endowment) and the National Science Foundation (IOS-2128266).

## Author contributions

ASG: Conceptualization, data curation, formal analysis, investigation, methodology, validation, writing original draft and review/editing

NCM: Conceptualization, data curation, investigation, formal analysis, methodology, writing original draft

SJK: Investigation, methodology JR: Investigation

HGB: Conceptualization, data curation, funding acquisition, investigation, methodology, project administration, resources, supervision, visualization, writing original draft and review/editing

## Acknowledgements

We thank past and present members of the Goodrich-Blair lab for feedback on the development of controls for bioinformatic co-occurrence analyses.

## Supplemental Files

**Supplemental File 1: T11SSGenomicCo-occurrence.xlsx**

Controlled co-occurrence analysis of all domains and genes significantly co-occurring with T11SS in a cluster of host-associated organisms (cluster 1) or a cluster of marine organisms (cluster 3). Significant genes were then clustered into UniRef50 clusters to combine close homologs and mined for predicted T11SS cargo proteins via homology to known cargo proteins.

**Supplemental File 2: Co-occurrenceFunctionalAnnotations.xlsx**

KEGG based analysis genes significantly co-occurring with host associated T11SS (cluster 1) and a marine T11SS (cluster 3).

**Supplemental File 3: Cluster3CargoLoci.xlsx**

Selected Cluster 3 predicted cargo proteins co-occurring with T11SS.

**Supplemental File 4: T11SS-DUF1194Co-occurrence.xlsx**

Co-occurrence of Marine associated T11SS alongside DUF1194 domain encoding genes, color coded to indicate host-associated organisms.

---

**Supplemental Figure 1.**
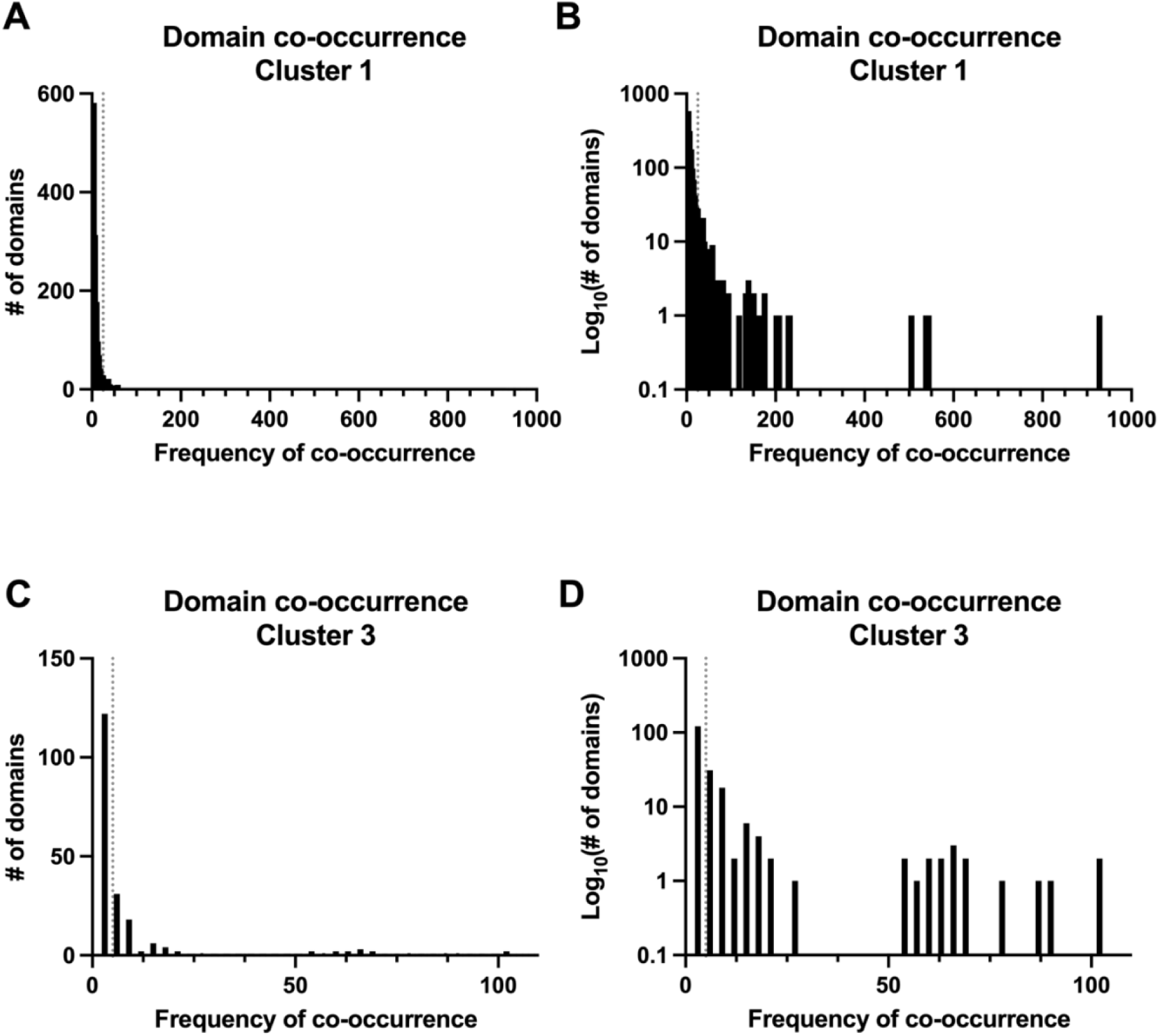
Histograms displaying the distribution of co-occurrence frequency amongst domains co-occurring with T11SS. Genomic neighborhood analysis detects many domains within assayed loci (6 open reading frames up- and down-stream of each T11SS), however the majority of domains only co-occur a small number of times. To filter out rare or spurious co-occurrences, a threshold of co-occurrence is established relative to the size of your dataset. Histograms depict the frequency of domain co-occurrence with animal-associated cluster 1 T11SS **(A)** and with marine-associated cluster 3 T11SS **(C)**. Log_10_ transformation of these values **(B, D)** helps to visualize domains with many co-occurrences. The dotted lines on all graphs show the thresholds chosen for filtering each dataset.

**Supplemental Figure 2.**
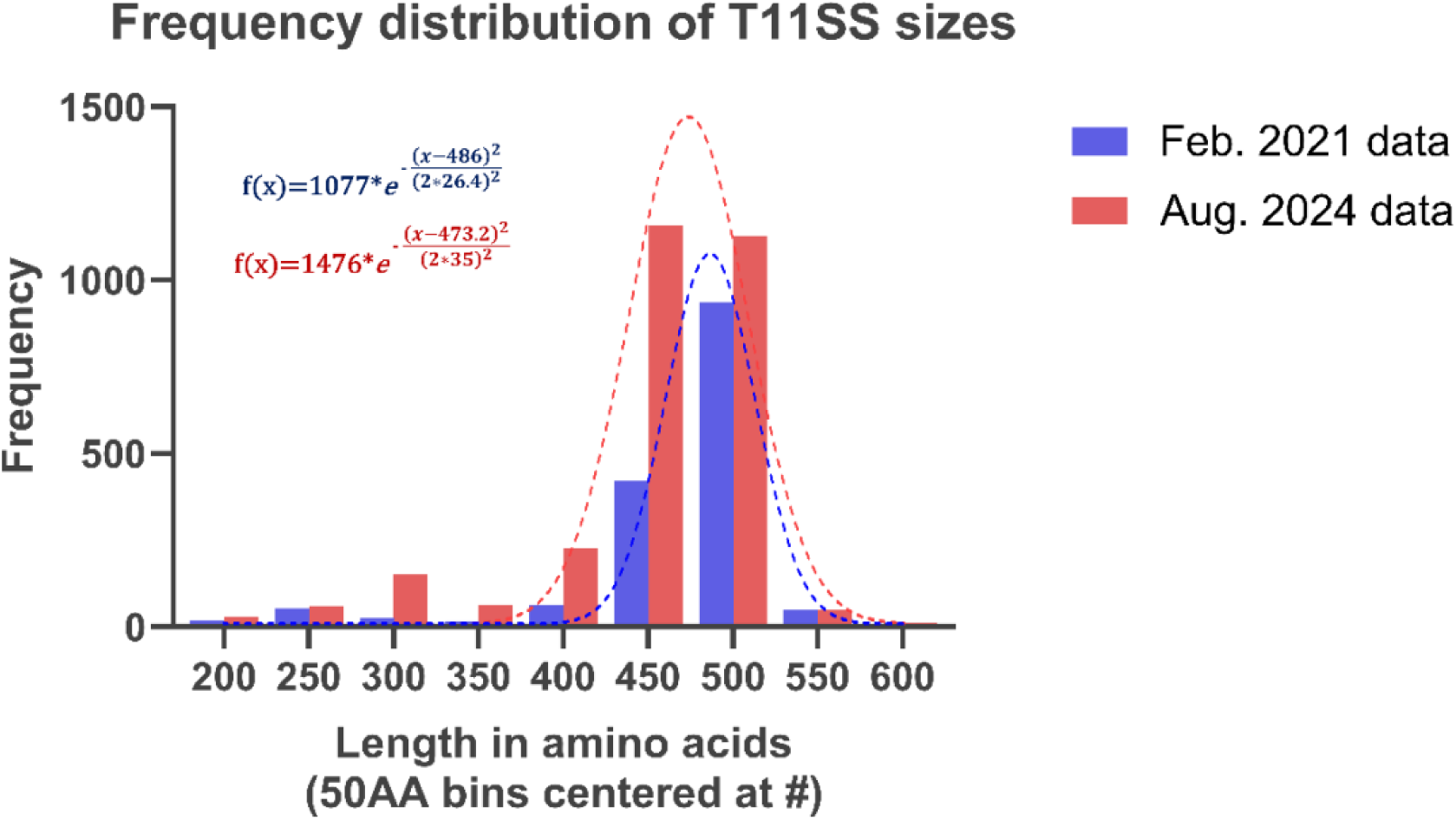
T11SS (DUF560) protein size distribution (in amino acids). After examination of the size distribution, median was chosen as an appropriate measure of representative protein size over mean due to left-skewed distribution. The blue dataset was used to perform the T11SS co-occurrence analysis described in this manuscript. In the intervening time more homologs have been added to the Pfam database, the red dataset reflects these additions and their impact on the size distribution.

**Supplemental Figure 3.**
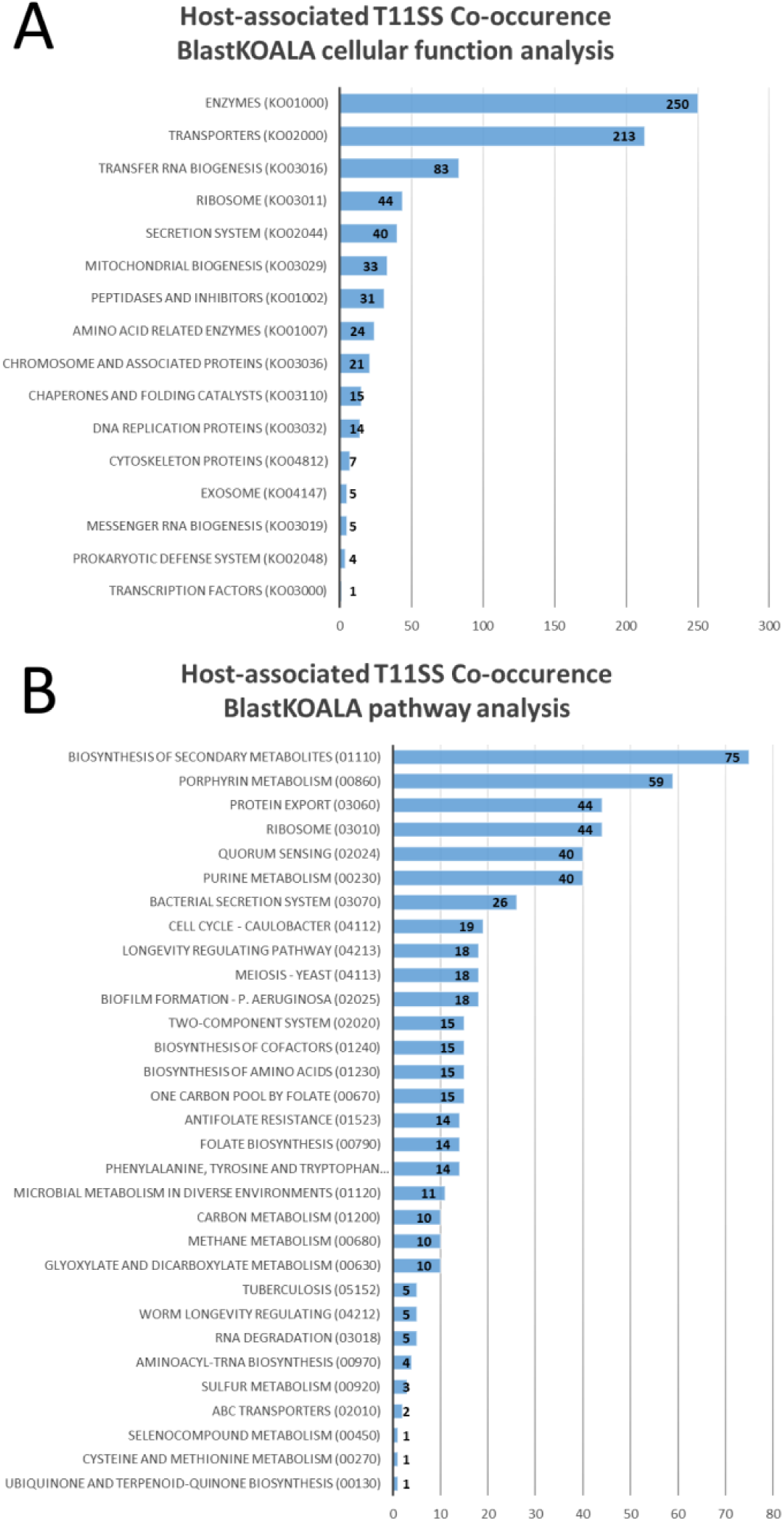
**BlastKOALA supports T11SS association with iron/heme uptake, protein export, and one-carbon metabolism**. BlastKOALA uses the BLAST alignment algorithm to assign functions to query sequences and reveal shared pathways. Cellular functions **(A)** were estimated using BRITE hierarchies and assigned, where possible, to known pathways **(B)** to detect commonalities.

**Supplemental Figure 4.**
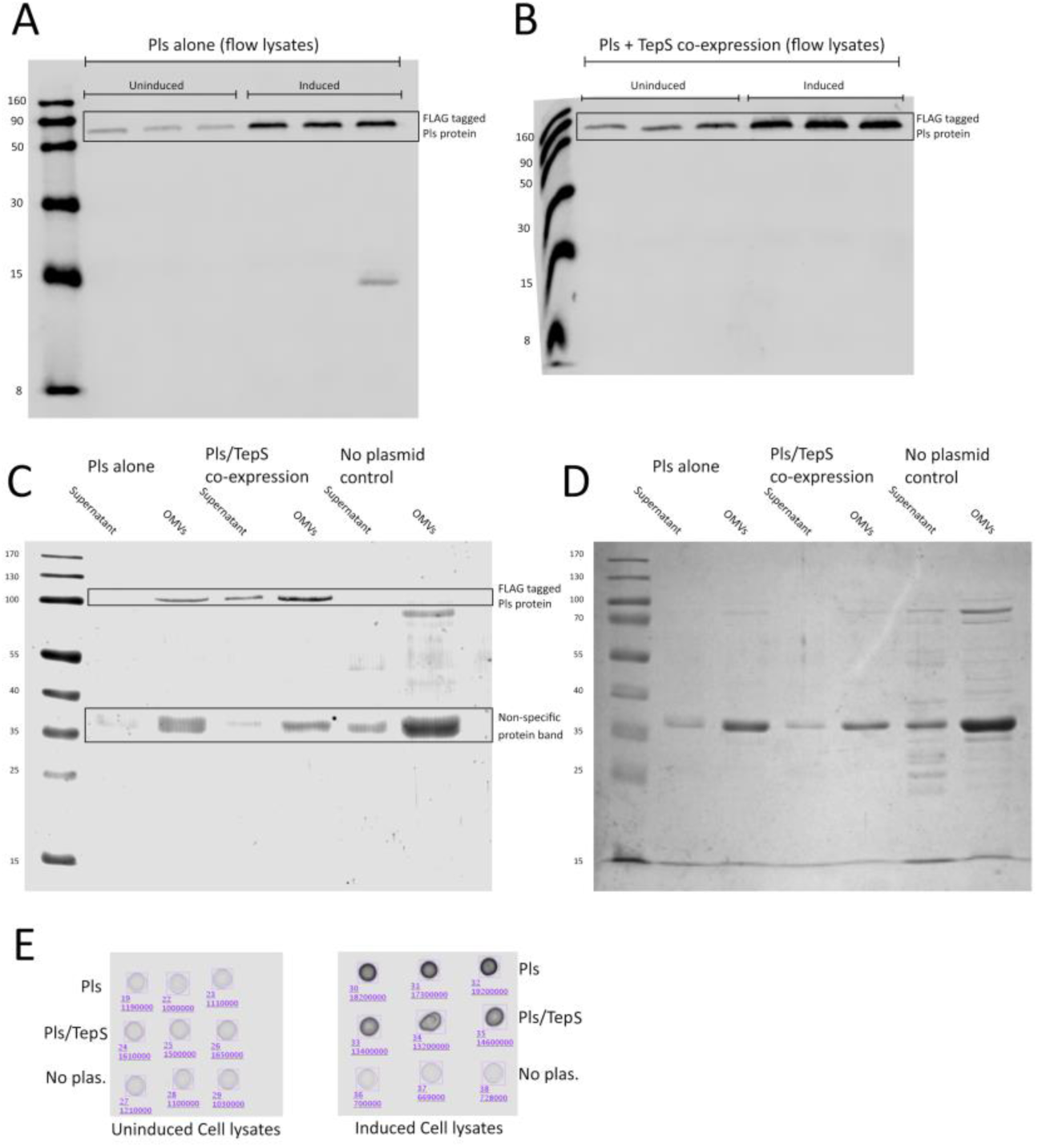
Localization of FLAG tagged Pls in the absence and presence of its cognate T11SS secretor. Samples from *E. coli* cells expressing Pls-FLAG alone or Pls-FLAG and its cognate T11SS (TepS) were probed with α-FLAG antibody using Western blotting (lysates from flow cytometry samples, supernatants, and extracellular vesicles) **(panels A-C)** or dot blots (lysates from supernatant collections) **(panel E)** to determine if expression of the T11SS-protein TepS impacted Pls protein localization. Representative Western blots demonstrating comparable levels of Pls expression in flow cytometry cultures in Pls only **(A)** and Pls and TepS **(B)** expressing cells. **(C)** A representative Western blot showing α-FLAG-antibody reactive supernatant proteins separated into soluble and vesicle fractions. **(D)** A representative Coomassie stain showing total supernatant protein content separated into soluble and vesicle fractions. **(E)** Immuno-dot blots demonstrating comparable levels of Pls expression in lysates from supernatant collection cultures.

**Supplemental Figure 5.**
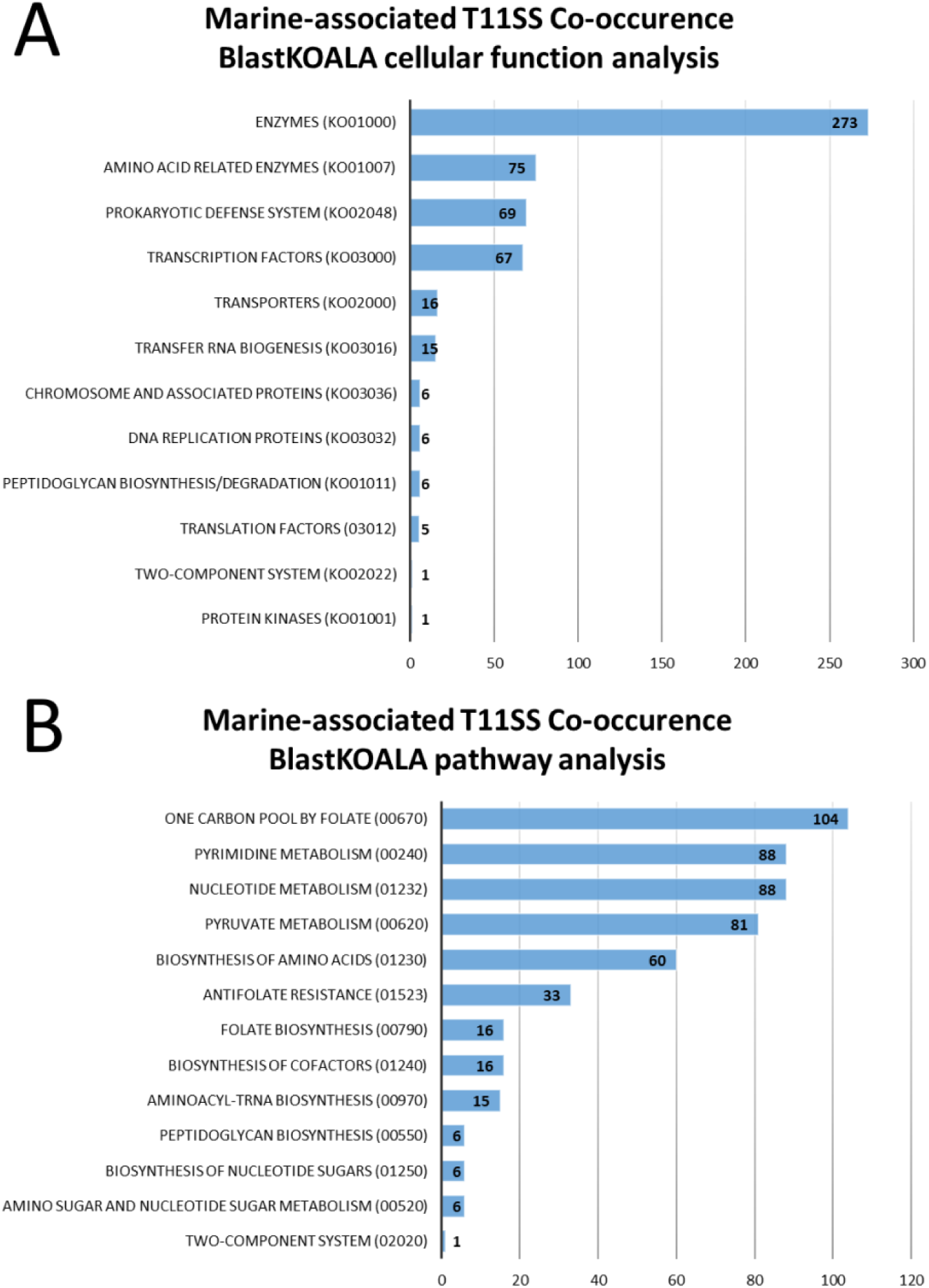
BlastKOALA supports T11SS association with one-carbon metabolism, nucleotide metabolism, and the glyoxalase-detoxification pathway. BlastKOALA uses the BLAST alignment algorithm to assign functions to query sequences and reveal shared pathways. Cellular functions (A) were estimated using BRITE hierarchies and assigned, where possible, to known pathways (B).

## Notes

https://doi.org/10.6084/m9.figshare.28514069.v2

### Competing Interest Statement

The authors have declared no competing interest.

### Summary of Updates

Additional data supporting the T11SS-dependent secretion of Pls; figshare links to supplementary data; revised figures and text for clarity.

## References

1. Grossman AS, Mucci NC, Kauffman SJ, Rafi J, Goodrich-Blair H. Bioinformatic discovery of type 11 secretion system (T11SS) cargo across the Proteobacteria supplemental data. Figshare. Online resource. Epub ahead of print 5 March 2025. DOI: 10.6084/m9.figshare.28514069.v2.

2. Mistry J, Chuguransky S, Williams L, Qureshi M, Salazar G, et al. Pfam: The protein families database in 2021. Nucleic Acids Res 2021;49:D412–D419.

3. Mudgal R, Sandhya S, Chandra N, Srinivasan N. De-DUFing the DUFs: Deciphering distant evolutionary relationships of domains of unknown function using sensitive homology detection methods. Biol Direct 2015;10:38.

4. Grossman AS, Mauer TJ, Forest KT, Goodrich-Blair H, Comstock L. A widespread bacterial secretion system with diverse substrates. mBio 2022;12:e01956–21.

5. Huynh MS, Hooda Y, Li YR, Jagielnicki M, Lai CC-L, et al. Reconstitution of surface lipoprotein translocation through the Slam translocon. Elife;11. Epub ahead of print April 2022. DOI: 10.7554/eLife.72822.

6. Shin HE, Pan C, Curran DM, Bateman TJ, Chong DHY, et al. Prevalence of Slam-dependent hemophilins in Gram-negative bacteria. J Bacteriol 2024;206:e00027–24.

7. Hooda Y, Lai CC-L, Moraes TF. Identification of a large family of Slam-dependent surface lipoproteins in Gram-negative bacteria. Front Cell Infect Microbiol;7. Epub ahead of print 2017. DOI: 10.3389/fcimb.2017.00207.

8. Hooda Y, Lai CC-L, Judd A, Buckwalter CM, Shin HE, et al. Slam is an outer membrane protein that is required for the surface display of lipidated virulence factors in *Neisseria*. Nat Microbiol;1. Epub ahead of print 2016. DOI: 10.1038/nmicrobiol.2016.9.

9. Cowles CE, Goodrich-Blair H. The *Xenorhabdus nematophila* nilABC genes confer the ability of *Xenorhabdus* spp. to colonize *Steinernema carpocapsae* nematodes. J Bacteriol 2008;190:4121– 4128.

10. Bateman TJ, Shah M, Ho TP, Shin HE, Pan C, et al. A Slam-dependent hemophore contributes to heme acquisition in the bacterial pathogen *Acinetobacter baumannii*. Nat Commun 2021;12:6270.

11. Latham RD, Torrado M, Atto B, Walshe JL, Wilson RK, et al. A heme-binding protein produced by *Haemophilus haemolyticus* inhibits non-typeable *Haemophilus influenzae*. Mol Microbiol 2020;113:381–398.

12. Grossman AS, Gell DA, Wu DG, Carper DL, Hettich RL, et al. Bacterial hemophilin homologs and their specific type eleven secretor proteins have conserved roles in heme capture and are diversifying as a family. J Bacteriol 2024;206:e00444–23.

13. Sam F, Brianna A, Arianna M, J HK, S RJ, et al. Heme sequestration by hemophilin from Haemophilus haemolyticus reduces respiratory tract colonization and infection with non-typeable Haemophilus influenzae. mSphere 2024;9:e00006–24.

14. Haft DH, Badretdin A, Coulouris G, DiCuccio M, Durkin AS, et al. RefSeq and the prokaryotic genome annotation pipeline in the age of metagenomes. Nucleic Acids Res 2024;52:D762–D769.

15. Rogozin IB, Makarova KS, Murvai J, Czabarka E, Wolf YI, et al. Connected gene neighborhoods in prokaryotic genomes. Nucleic Acids Res 2002;30:2212–2223.

16. Mahdavi MA, Lin Y-H. False positive reduction in protein-protein interaction predictions using gene ontology annotations. BMC Bioinformatics 2007;8:262.

17. Tietz JI, Schwalen CJ, Patel PS, Maxson T, Blair PM, et al. A new genome-mining tool redefines the lasso peptide biosynthetic landscape. Nat Chem Biol 2017;13:470–478.

18. The UniProt Consortium. UniProt: The universal protein knowledge base in 2021. Nucleic Acids Res 2021;49:D480–D489.

19. Chen I-MA, Chu K, Palaniappan K, Ratner A, Huang J, et al. The IMG/M data management and analysis system v.7: content updates and new features. Nucleic Acids Res 2023;51:D723–D732.

20. Mukherjee S, Stamatis D, Li CT, Ovchinnikova G, Bertsch J, et al. Twenty-five years of Genomes OnLine Database (GOLD): data updates and new features in v.9. Nucleic Acids Res 2023;51:D957–D963.

21. Jumper J, Evans R, Pritzel A, Green T, Figurnov M, et al. Highly accurate protein structure prediction with AlphaFold. Nature 2021;596:583–589.

22. Mirdita M, Schütze K, Moriwaki Y, Heo L, Ovchinnikov S, et al. ColabFold: Making protein folding accessible to all. Nat Methods 2022;19:679–682.

23. Abramson J, Adler J, Dunger J, Evans R, Green T, et al. Accurate structure prediction of biomolecular interactions with AlphaFold 3. Nature 2024;630:493–500.

24. Aramaki T, Blanc-Mathieu R, Endo H, Ohkubo K, Kanehisa M, et al. KofamKOALA: KEGG Ortholog assignment based on profile HMM and adaptive score threshold. Bioinformatics 2020;36:2251–2252.

25. Kanehisa M, Sato Y, Morishima K. BlastKOALA and GhostKOALA: KEGG tools for functional characterization of genome and metagenome sequences. J Mol Biol 2016;428:726–731.

26. Suzek BE, Huang H, McGarvey P, Mazumder R, Wu CH. UniRef: comprehensive and non-redundant UniProt reference clusters. Bioinformatics 2007;23:1282–1288.

27. Buchan DWA, Jones DT. The PSIPRED protein analysis workbench: 20 years on. Nucleic Acids Res 2019;47:W402–W407.

28. Bhasin A, Chaston JM, Goodrich-Blair H. Mutational analyses reveal overall topology and functional regions of NilB, a bacterial outer membrane protein required for host association in a model of animal-microbe mutualism. J Bacteriol 2012;194:1763–1776.

29. Miroux B, Walker JE. Over-production of proteins in *Escherichia coli*: mutant hosts that allow synthesis of some membrane proteins and globular proteins at high levels. J Mol Biol 1996;260:289–298.

30. Dumon-Seignovert L, Cariot G, Vuillard L. The toxicity of recombinant proteins in *Escherichia coli*: a comparison of overexpression in BL21(DE3), C41(DE3), and C43(DE3). Protein Expr Purif 2004;37:203–206.

31. Orchard SS, Goodrich-Blair H. Identification and functional characterization of a *Xenorhabdus nematophila* oligopeptide permease. Appl Environ Microbiol 2004;70:5621–5627.

32. Grossman AS, Escobar CA, Mans EJ, Mucci NC, Mauer TJ, et al. A surface exposed, two-domain lipoprotein cargo of a type XI secretion system promotes colonization of host intestinal epithelia expressing glycans. Frontiers in Microbiology;13. https://www.frontiersin.org/article/10.3389/fmicb.2022.800366 (2022).

33. Koontz L. Chapter one - TCA precipitation. In: Lorsch JBT-M in E (editor). Laboratory methods in enzymology. Academic Press. pp. 3–10.

34. Tukey JW. Comparing individual means in the analysis of variance. Biometrics 1949;5:99–114.

35. Kanehisa M, Furumichi M, Tanabe M, Sato Y, Morishima K. KEGG: New perspectives on genomes, pathways, diseases and drugs. Nucleic Acids Res 2017;45:D353–D361.

36. Dobrindt U, Reidl J. Pathogenicity islands and phage conversion: Evolutionary aspects of bacterial pathogenesis. International Journal of Medical Microbiology 2000;290:519–527.

37. Almagro Armenteros JJ, Tsirigos KD, Sønderby CK, Petersen TN, Winther O, et al. SignalP 5.0 improves signal peptide predictions using deep neural networks. Nat Biotechnol 2019;37:420– 423.

38. Teufel F, Almagro Armenteros JJ, Johansen AR, Gíslason MH, Pihl SI, et al. SignalP 6.0 predicts all five types of signal peptides using protein language models. Nat Biotechnol 2022;40:1023– 1025.

39. Skaf MS, Polikarpov I, Stanković IM. A linker of the proline-threonine repeating motif sequence is bimodal. J Mol Model 2020;26:178.

40. Swearingen KE, Lindner SE, Shi L, Shears MJ, Harupa A, et al. Interrogating the *Plasmodium* sporozoite surface: identification of surface-exposed proteins and demonstration of glycosylation on CSP and TRAP by mass spectrometry-based proteomics. PLoS Pathog 2016;12:e1005606.

41. Sieber CMK, Paul BG, Castelle CJ, Hu P, Tringe SG, et al. Unusual metabolism and hypervariation in the genome of a Gracilibacterium (BD1-5) from an oil-degrading community. mBio;10. Epub ahead of print November 2019. DOI: 10.1128/mBio.02128-19.

42. Calmettes C, Alcantara J, Yu R-H, Schryvers AB, Moraes TF. The structural basis of transferrin sequestration by transferrin-binding protein B. Nat Struct Mol Biol 2012;19:358–360.

43. Ostberg K, DeRocco A, Mistry S, Dickinson Mary K, Cornelissen CN. Conserved regions of gonococcal TbpB are critical for surface exposure and transferrin iron utilization. Infect Immun 2013;81:3442–3450.

44. Bleiziffer I, Eikmeier J, Pohlentz G, McAulay K, Xia G, et al. The Plasmin-sensitive protein Pls in methicillin-resistant *Staphylococcus aureus* (MRSA) is a glycoprotein. PLoS Pathog 2017;13:e1006110.

45. Savolainen K, Paulin L, Westerlund-Wikström B, Foster TJ, Korhonen TK, et al. Expression of *pls*, a gene closely associated with the *mecA* gene of methicillin-resistant *Staphylococcus aureus*, prevents bacterial adhesion *in vitro*. Infect Immun 2001;69:3013–3020.

46. Liang KYH, Orata FD, Boucher YF, Case RJ. Roseobacters in a sea of poly- and paraphyly: Whole genome-based taxonomy of the family Rhodobacteraceae and the proposal for the split of the “Roseobacter Clade” into a novel family, Roseobacteraceae fam. nov. Front Microbiol;12. Epub ahead of print 2021. DOI: 10.3389/fmicb.2021.683109.

47. Crenn K, Serpin D, Lepleux C, Overmann J, Jeanthon C. *Silicimonas algicola* gen. nov., sp. nov., a member of the Roseobacter clade isolated from the cell surface of the marine diatom Thalassiosira delicatula. Int J Syst Evol Microbiol 2016;66:4580–4588.

48. Kessner L, Spinard E, Gomez-Chiarri M, Rowley DC, Nelson DR. Draft genome sequence of *Aliiroseovarius crassostreae* CV919-312, the causative agent of Roseovarius Oyster Disease (formerly Juvenile Oyster Disease). Genome Announc;4. Epub ahead of print March 2016. DOI: 10.1128/genomeA.00148-16.

49. Kim Y-O, Park S, Nam B-H, Lee C, Park J-M, et al. *Ascidiaceihabitans donghaensis* gen. nov., sp. nov., isolated from the golden sea squirt Halocynthia aurantium. Int J Syst Evol Microbiol 2014;64:3970–3975.

50. Ivanova EP, Gorshkova NM, Sawabe T, Zhukova N V, Hayashi K, et al. *Sulfitobacter delicatus* sp. nov. and *Sulfitobacter dubius* sp. nov., respectively from a starfish (*Stellaster equestris*) and sea grass (*Zostera marina*). Int J Syst Evol Microbiol 2004;54:475–480.

51. Chen M-H, Sheu S-Y, Chen CA, Wang J-T, Chen W-M. *Roseivivax isoporae* sp. nov., isolated from a reef-building coral, and emended description of the genus *Roseivivax*. Int J Syst Evol Microbiol 2012;62:1259–1264.

52. Alamos P, Tello M, Bustamante P, Gutiérrez F, Shmaryahu A, et al. Functionality of tRNAs encoded in a mobile genetic element from an acidophilic bacterium. RNA Biol 2018;15:518–527.

53. Baek M, DiMaio F, Anishchenko I, Dauparas J, Ovchinnikov S, et al. Accurate prediction of protein structures and interactions using a three-track neural network. Science (1979) 2021;373:871–876.

54. Bergelson JM, Hemler ME. Integrin-Ligand Binding: Do integrins use a ‘MIDAS touch’ to grasp an Asp? Current Biology 1995;5:615–617.

55. Cantí C, Nieto-Rostro M, Foucault I, Heblich F, Wratten J, et al. The metal-ion-dependent adhesion site in the Von Willebrand factor-A domain of alpha2delta subunits is key to trafficking voltage-gated Ca2+ channels. Proc Natl Acad Sci U S A 2005;102:11230–11235.

56. Esch R, Merkl R. Conserved genomic neighborhood is a strong but no perfect indicator for a direct interaction of microbial gene products. BMC Bioinformatics 2020;21:5.

57. Hantke K. Is the bacterial ferrous iron transporter FeoB a living fossil? Trends Microbiol 2003;11:192–195.

58. Ollis AA, Postle K. ExbD mutants define initial stages in TonB energization. J Mol Biol 2012;415:237–247.

59. Levengood SK, Beyer WFJ, Webster RE. TolA: a membrane protein involved in colicin uptake contains an extended helical region. Proc Natl Acad Sci U S A 1991;88:5939–5943.

60. Tjalsma H, Kontinen VP, Prágai Z, Wu H, Meima R, et al. The role of lipoprotein processing by signal peptidase II in the Gram-positive eubacterium *Bacillus subtilis*. Signal peptidase II is required for the efficient secretion of alpha-amylase, a non-lipoprotein. J Biol Chem 1999;274:1698–1707.

61. Agrawal N, Lesley SA, Kuhn P, Kohen A. Mechanistic studies of a flavin-dependent thymidylate synthase. Biochemistry 2004;43:10295–10301.

62. Dawadi S, Kordus SL, Baughn AD, Aldrich CC. Synthesis and analysis of bacterial folate metabolism intermediates and antifolates. Org Lett 2017;19:5220–5223.

63. Luk LYP, Javier Ruiz-Pernía J, Dawson WM, Roca M, Loveridge EJ, et al. Unraveling the role of protein dynamics in dihydrofolate reductase catalysis. Proc Natl Acad Sci U S A 2013;110:16344– 16349.

64. Mendel RR. The molybdenum cofactor. J Biol Chem 2013;288:13165–13172.

65. Zhong Q, Kobe B, Kappler U. Molybdenum enzymes and how they support virulence in pathogenic bacteria. Frontiers in Microbiology;11. https://www.frontiersin.org/articles/10.3389/fmicb.2020.615860 (2020).

66. Van Bibber M, Bradbeer C, Clark N, Roth JR. A new class of cobalamin transport mutants (*btuF*) provides genetic evidence for a periplasmic binding protein in *Salmonella typhimurium*. J Bacteriol 1999;181:5539–5541.

67. Peyvandi F, Garagiola I, Baronciani L. Role of von Willebrand factor in the haemostasis. Blood Transfus 2011;9 Suppl 2:s3–8.

68. Von Willebrand EA. Hereditary pseudohaemophilia. Haemophilia 1999;5:223–31; discussion 222.

69. Islam ST, Jolivet NY, Cuzin C, Belgrave AM, My L, et al. Unmasking of the von Willebrand A-domain surface adhesin CglB at bacterial focal adhesions mediates myxobacterial gliding motility. Sci Adv 2023;9:eabq0619.

70. Aravind L, Iyer LM, Burroughs AM. Discovering biological conflict systems through genome analysis: Evolutionary principles and biochemical novelty. Annu Rev Biomed Data Sci 2022;5:367– 391.

71. Kaur G, Burroughs AM, Iyer LM, Aravind L. Highly regulated, diversifying NTP-dependent biological conflict systems with implications for the emergence of multicellularity. Elife 2020;9:e52696.

72. Lee C, Park C. Bacterial responses to glyoxal and methylglyoxal: Reactive electrophilic species. Int J Mol Sci;18. Epub ahead of print January 2017. DOI: 10.3390/ijms18010169.

73. Anaya-Sanchez A, Feng Y, Berude JC, Portnoy DA. Detoxification of methylglyoxal by the glyoxalase system is required for glutathione availability and virulence activation in *Listeria monocytogenes*. PLoS Pathog 2021;17:e1009819.

74. Fantappiè L, Irene C, De Santis M, Armini A, Gagliardi A, et al. Some Gram-negative lipoproteins keep their surface topology when transplanted from one species to another and deliver foreign polypeptides to the bacterial surface. Molecular & Cellular Proteomics 2017;16:1348 LP – 1364.

75. Konar M, Rossi R, Walter H, Pajon R, Beernink PT. A mutant library approach to identify improved meningococcal Factor H binding protein vaccine antigens. PLoS One 2015;10:e0128185.

76. Cao Y, Gao L, Zhang L, Zhou L, Yang J, et al. Genome-wide screening of lipoproteins in *Actinobacillus pleuropneumoniae* identifies three antigens that confer protection against virulent challenge. Sci Rep 2020;10:2343.

77. Schmidt MA, Riley LW, Benz I. Sweet new world: Glycoproteins in bacterial pathogens. Trends Microbiol 2003;11:554–561.

78. Tortorelli G, Rautengarten C, Bacic A, Segal G, Ebert B, et al. Cell surface carbohydrates of symbiotic dinoflagellates and their role in the establishment of cnidarian-dinoflagellate symbiosis. ISME J 2022;16:190–199.

79. Zhou M, Wu H. Glycosylation and biogenesis of a family of serine-rich bacterial adhesins. Microbiology (N Y) 2009;155:317–327.

